# Talin regulates steady-state tensional homeostasis to drive vascular morphodynamics and cancer

**DOI:** 10.1101/2022.08.03.502607

**Authors:** Pinelopi Nikolopoulou, Christina Arapatzi, Georgia Rouni, Demosthenis Mitrossilis, Anastasios Gaitanis, Constantinos D. Anagnostopoulos, Sofia Grammenoudi, Vassiliki Kostourou

## Abstract

The mechanical properties of the extracellular environment emerge as critical regulators of cellular functions. Cell mechanotransduction is mainly studied *in vitro* at initial stages of cell adhesion and very little is known about the mechanoresponses of cells with established tensional dynamics, resembling cells embedded in tissues. Here, we provide *in vivo* evidence that talin-dependent cell-matrix adhesions are global regulators of vascular mechanics and establish talin as an essential and required mechanosensor in neovessels and already developed tumours. At the molecular level, we demonstrate that talin exploits alternative mechanisms to dynamically-adjust the mechanical integrity of endothelial cells. Our mutational studies indicate a previously unknown role for the requirement of the talin-head in mechanosensing and demonstrate that the talin-head and the talin-rod alone are sufficient to maintain mechanical stability of endothelial cells. Overall, our results underpin the significance of mechanical signals in regulating vascular morphology in steady-state conditions and ultimately modulate cancer progression.

**Talin mechanosensing is required to maintain cell morphology and control developmental and tumour angiogenesis.**

## Introduction

In the tumour microenvironment both cancer cells and tumour-stroma communicate via diverse cues, including growth factors, cytokines, cell-cell signalling and components of extracellular matrix (**ECM**), to drive excessive and abnormal vascular function and tumour development (*1, 2*). Besides these biochemical signals, the mechanical properties of the tumour microenvironment critically determine tumour progression and response to treatment (*3*). How endothelial cells (**ECs**) that compose blood vessels sense their microenvironment to move, change shape and proliferate remains poorly understood.

Cell-matrix adhesions are major sites for the integration of chemical and mechanical signals from the microenvironment (*4*-*7*). They comprise a complex and highly dynamic network of adaptor and signalling proteins, termed the adhesome, which is formed around integrin transmembrane receptors that connect the ECM to the cell cytoskeleton (*8*-*11*). The function of the adhesome is to support physical adhesion and co-ordinate cell differentiation, migration, proliferation and survival, (*12*-*14*)all processes essential for vascular morphogenesis. Very little is known about the *in vivo* function of adhesome members in tumour vasculature (*15*) and even less is understood about their involvement in sensing and responding to the mechanical properties of tumours (*16*-*19*).

Talin is a central adhesome scaffolding protein with established roles in integrin activation and adhesion formation and maturation (*20*-*24*). Talin is composed by N-terminus head domain consisting of a non-canonical FERM domain, a flexible linker and a rod domain comprising 13 α-helical bundles (subdomains R1-R13), followed by the C-terminal dimerization domain (DD) (*25*). Talin directly binds to the cytoplasmic tails of β-integrins and to actin filaments creating a minimal physical linkage between ECM and the cytoskeleton. It also contains binding sites for other adhesome members and membrane phospholipids (PIP3) (*23, 26*). Accumulating *in vitro* data points to an important function for talin in cell mechanotransduction (*27, 28*). Biophysical studies show that the a-helical bundles of the talin rod unfold in response to force, indicating that talin can sense and bear forces at cell-matrix adhesions (*29*-*35*). Traction force experiments in cells deficient in talin isoforms or cells expressing various talin-rod domain mutants confirm that talin is required for force transmission during cell spreading and initial development of cell-matrix adhesions (*21, 36*-*42*). Although these studies provide important insight into the role of talin in force transmission during early events of cell attachment and adhesion formation, they do not address whether and how talin affects mechanoresponses in well-spread cells with established adhesions in tissues *in vivo*. Therefore, a gap remains in translating these molecular and cell-based findings to the whole organism and to pathological situations such as cancer.

Given that endothelial talin expression is essential for embryonic angiogenesis (*43*) and vascular homeostasis (*44*), we set out to determine the molecular mechanism of talin’s action in steady-state EC dynamics in newly formed vessels and established tumours. We found that inducible deletion of endothelial talin at different points of tumour development ceased tumour growth and caused tumour regression by disrupting EC morphodynamics and disintegrating the newly formed vascular network. Taking advantage of our inducible system of talin deletion, we demonstrate that talin is required for steady-state EC rigidity-triggered responses that control cell morphology. In contrast to initial cell spreading events, we found that either the talin-head or the talin-rod alone can transmit forces and balance cellular tension in ECs, thus revealing a previously undiscovered mechanical function for the talin head. We propose that talin exploits partially redundant mechanisms to stabilise cell morphology and elicit rigidity-triggered responses in blood vessels. Our data establish a new role for talin mechanosensing in maintaining force equilibrium in well-attached cells, which resemble *in vivo* conditions. Additionally, our *in vivo* studies link previous *in vitro* evidence of talin’s mechanical role to a functional outcome and a pathological condition in a mammalian organism. Overall, our data expand our knowledge of how endothelial cell-matrix adhesions regulate blood vessel morphodynamics and reveal the fundamental contribution of mechanical cues in vascular function and tumour growth, paving the way for novel therapeutic interventions.

## Results

### Endothelial talin deletion limits tumour growth and causes tumour regression by inhibiting blood vessel morphodynamics

Despite the essential role of EC in tumour growth, how endothelial adhesome proteins affect cancer remains poorly understood. Hence, we set out to examine the impact of endothelial talin deletion at different stages of tumour development, using tamoxifen inducible *Pdgfb-*iCreER^T2^; Talin1^fl/fl^ mice (*45*-*47*). Mice were subcutaneously injected with syngeneic mouse B16F0 melanoma or CMT19T adenocarcinoma cells, and EC-specific talin deletion was induced by tamoxifen administration. We verified the *in vivo* elimination of talin expression by western blots of isolated lung ECs derived from tumour-bearing tamoxifen-treated (EC-Talin^KO^) and *Pdgfb-*iCreER^T2^ negative; Talin1^fl/fl^ (EC-Talin^WT^) mice (Fig. S1). Induction of EC talin deletion on the same day as tumour cell inoculation completely abolished tumour development (Fig.1A). Tamoxifen administration after tumour initiation also resulted in much smaller tumours in EC-Talin^KO^ mice compared to EC-Talin^WT^ littermates (Fig. S2A). Strikingly, deletion of EC talin in established tumours, dramatically reduced tumour volume in EC-Talin^KO^ mice compared to EC-Talin^WT^ littermates (Fig. 1A), not only arresting tumour growth but also promoting tumour shrinkage (Fig. S2B). To further investigate whether loss of EC talin causes tumour regression, we performed micro-PET/CT imaging on established B16F0 melanomas before and after induction of EC talin deletion (Fig. 1B). Tumours grew at similar rates before tamoxifen administration on day 10 post-inoculation, as indicated by the mean standardized uptake value (SUV) for 2-^18^F-fluoro-2-deoxy-D-glucose (^18^F-FDG), and the total lesion glycolysis (TGL) index, both measurements used in a clinical setting to evaluate tumour responses to treatment. Both SUV and TGL values were significantly reduced upon loss of EC talin, demonstrating reduction in viable tumour mass (Fig. 1B). Consistent with these findings, EC talin deletion in established tumours significantly reduced proliferation of tumour cells as determined by quantification of Ki67 immunostaining, and increased tumour cell apoptosis, as indicated by quantification of activated caspace-3 positive tumour area (Fig. 1C).

**Fig. 1.**
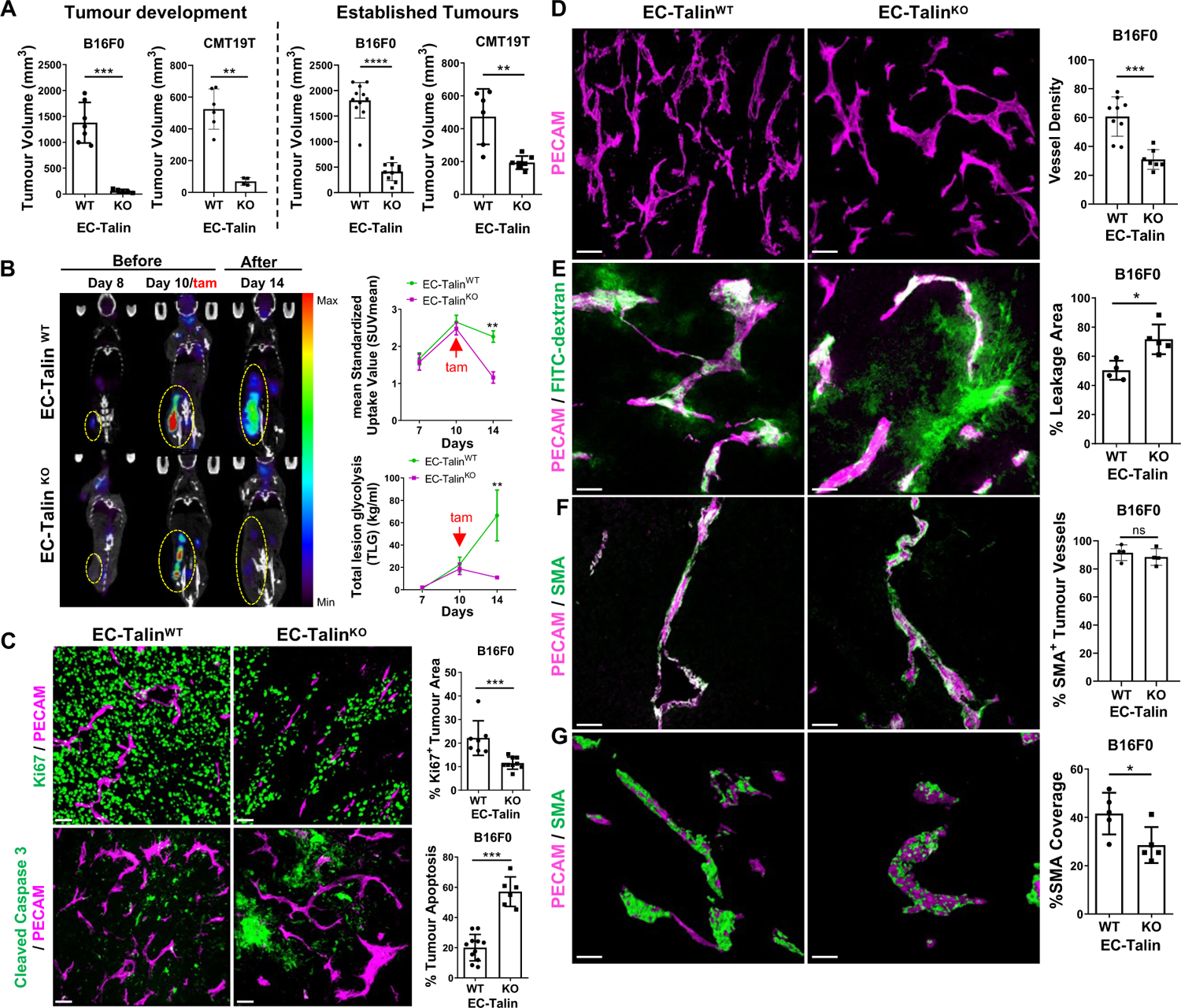
Endothelial talin deletion diminishes tumour progression and disrupts tumour blood vessel morphogenesis and function. (**A**) Tumour volume of subcutaneous B16F0 and CMT19T tumours grown in EC-Talin^KO^ mice and EC-Talin^WT^ littermates for 14 days and 16 days respectively. EC-Talin deletion occurred the same day as tumour inoculation (tumour development data) and 10 days after tumour inoculation (established tumour data). For tumour development studies: B16F0, n=7 EC-Talin^WT^ and 5 EC-Talin^KO^, **P=0.0025; CMT19T, n=6 EC-Talin^WT^ and 4 EC-Talin^KO^, **P=0.0095. For established tumour studies: B16F0, n=11 EC-Talin^WT^ and 10 EC-Talin^KO^, ****P<0.0001; CMT19T, n=6 EC-Talin^WT^ and 7 EC-Talin^KO^, **P=0.0023. (**B**) Whole-body micro-PET/CT images (coronal view), after intravenous injection of ^18^F-FDG in mice bearing B16F0 tumours, before (day 7, 10) and after endothelial talin deletion (day 14). Graphs represent the mean Standard Uptake Value (SUV) and the total glycolysis rate (TLG) between EC-Talin^KO^ and EC-Talin^WT^ littermates; n=7 mice per genotype, day 7 and day 10: ns, no statistically significant difference; day 14: **P=0.0023 (SUV analysis), **P=0.076 (TLG analysis). PET images of tumours were colour-coded on the same scale. (**C**) Representative images of tumour sections from EC-Talin^WT^ and EC-Talin^KO^ mice stained with antibodies against PECAM and Ki67 (upper panels) and PECAM and cleaved caspase 3 (bottom panel). Bar charts represent the percentage of Ki-67 and cleaved caspase 3 positive tumour area over the total tumour area, respectively; for Ki67 analysis: n=7 EC-Talin^WT^ and 9 EC-Talin^KO^ B16F0 tumours, ***P=0.0002; for cleaved caspase 3 analysis: n=11 EC-Talin^WT^ and 6 EC-Talin^KO^ B16F0 tumours, ***P=0.0002. (**D**) Confocal images of 50mm tumour sections from EC-Talin^WT^ and EC-Talin^KO^ mice stained with antibodies against PECAM to demarcate blood vessels. Bar chart presents blood vessel density as the number of blood vessels per mm^2^ of midline tumour area; n=11 EC-Talin^WT^ and 6 EC-Talin^KO^, ***P=0.0007. (**E**) Confocal images of 50mm B16F0 tumour sections showing FITC-dextran leakiness from PECAM-stained vessels. Bar chart represents the percentage of extravascular FITC-Dextran area over the total FITC-Dextran area n=4 EC-Talin^WT^ and 5 EC-Talin^KO^, *P=0.0159. (**F**) Confocal images of thin B16F0 tumour sections stained for pericytes with aSMA and blood vessels with PECAM. Bar chart represents the percentage of blood vessels positive for aSMA staining divided by the total number of vessels; n=4 EC-Talin^WT^ and 4 EC-Talin^KO^ B16F0 tumours, ns. (**G**) Representative images of the aSMA- and PECAM-staining surface areas, denoting the direct physical contact between pericytes and the endothelium in 50mm midline tumour sections. Bar chart represents the percentage of PECAM staining area that overlaps with SMA staining area divided by the total PECAM staining area; n=5 EC-Talin^WT^ and 5 EC-Talin^KO^ B16F0 tumours, *P=0.0317. Scale bars, 50 µm. All bar charts represent the mean value ± s.e.m.; Statistical analysis, Mann–Whitney rank-sum test.

Examination of tumour vasculature in established tumours revealed that EC loss of talin reduced tumour blood vessel density in both B16F0 melanoma and CMT19T adenocarcinoma, detected by immunostaining of tumour sections for PECAM (Fig. 1D and Fig. S2C). The percentage of perfused tumour blood vessels was significantly lower in mice with EC-talin deletion compared to wild type (Fig. S2D), and vascular integrity was disrupted as shown by the increased leakiness of intravenously injected FITC-Dextran in EC-Talin^KO^ mice compared to in EC-Talin^WT^ littermates (Fig. 1E). These vascular defects were accompanied by increased hypoxia (Fig. S3A) without any differences in macrophage tumour infiltration, as shown by the quantification of CD11b (Fig. S3B) and F4/80 (Fig. S3C) immunostaining of tumour sections. Moreover, pericytes were loosely attached in blood vessels lacking EC talin. Although there was no difference in the percentage of aSMA-positive blood vessels in tumours grown in EC-Talin^KO^ and EC-Talin^WT^ mice (Fig. 1F), the surface attachment area of smooth muscle cells with ECs was significantly reduced (Fig. 1G). Taken together, these data demonstrate that EC-talin is essential for tumour vascular formation and function to drive cancer development.

### Loss of endothelial talin distorts vascular network morphology by creating viable cell aggregates

To decipher whether the defective tumour vasculature in mice lacking endothelial talin is because of impaired formation or maintenance of the newly formed vascular network, we performed the *ex vivo* aortic ring assay. We isolated aortic rings from *Pdgfb-*iCreER^T2^; Talin1^fl/fl^ mice and deleted endothelial talin by 4-OH tamoxifen (4-OHT) treatment at two different time-points of angiogenic sprouting. Similar to tumours, when we deleted endothelial talin at the initial stages of vessel outgrowth, no sprouts were forming (Fig. 2A). Deletion of endothelial talin when initial sprouts had developed, severely reduced vessel branching, ceased vessel expansion and ruptured the newly formed spouts (Fig. 2A, movie S1 and S2).

**Fig. 2.**
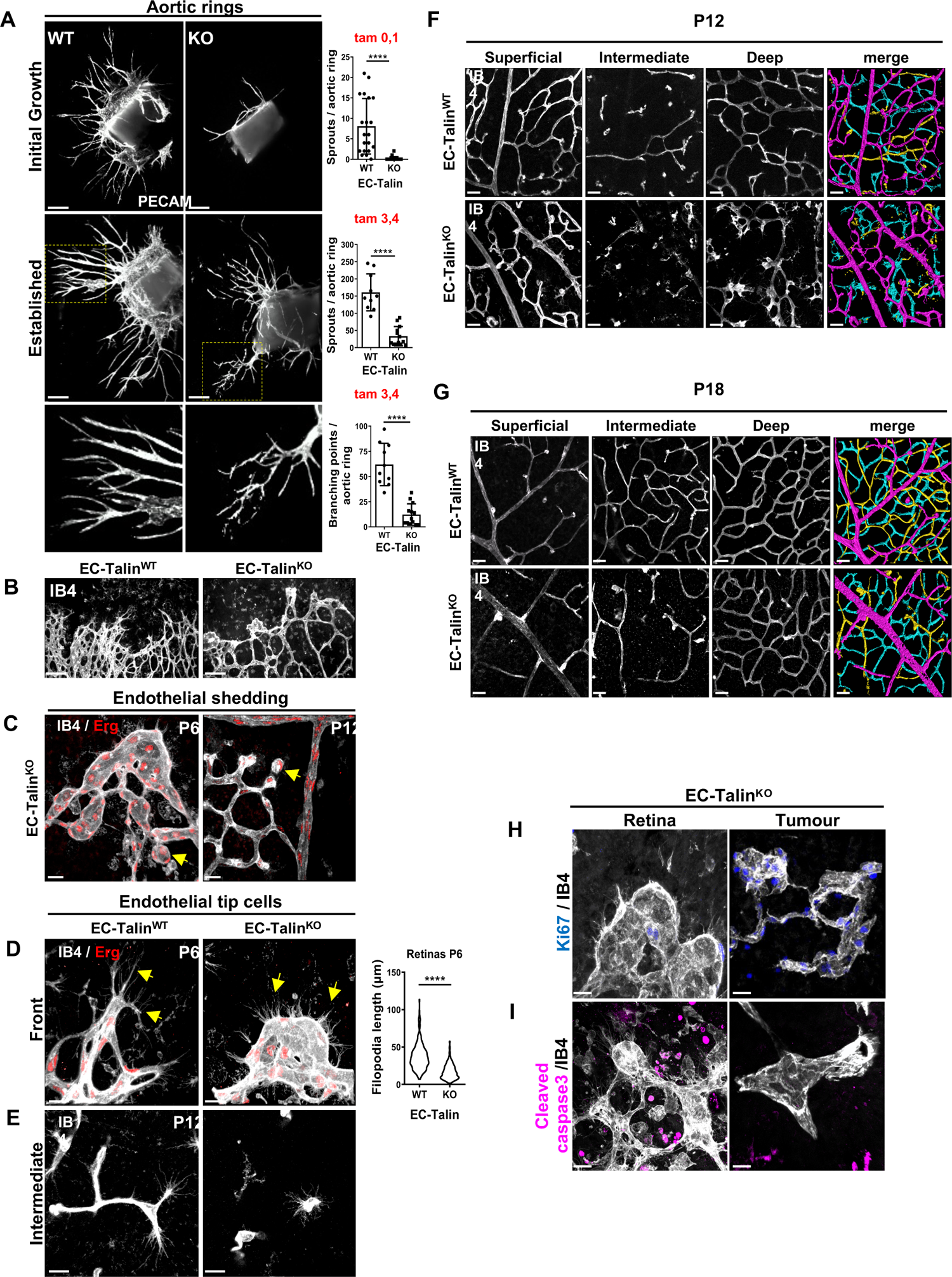
Loss of endothelial Talin causes severe vascular malformations and eliminates growth and remodeling of blood vessels. (**A**) Images of PECAM-stained aortic ring sprouts. Talin deletion is induced before the development of sprouts (initial growth) and in established sprouts. Insets illustrate the disrupted sprout network upon talin deletion in established sprouts. Bar charts represent the number of sprouts per aortic ring, during the two time-points of talin deletion and the number of branching points per aortic ring upon talin deletion in established sprouts, mean ± s.e.m.; for initial sprouts: n=20 EC-Talin^WT^ and 21 EC-Talin^KO^ aortic rings, ****P<0.0001; for established sprouts: n=10 wt and 13 KO aortic rings, ****P<0.0001. (**B**) Flat-mounted P6 mouse retina stained with isolectin-B4 (IB4), showing reduced vessel density and blunted angiogenic front in EC-TalinKO mice compared to EC-Talin^WT^ littermates. (**C**) EC-Talin^KO^ P6 and P12 retinas labeled with IB4 (white) and Erg (red) to mark ECs nuclei indicate endothelial shedding (yellow arrows). (**D**) EC-Talin deficiency resulted in shorter filopodia, indicated by yellow arrows, at vascular front. Violin plots display the length of individual filopodia per genotype; n=7 EC-Talin^WT^ and 9 EC-Talin^KO^, ****P<0.0001. (**E**) Representative confocal images of tip cells in the intermediate layer showing abnormal filopodia extensions and impaired invasion in EC-Talin^KO^ P12 retinas compared to EC-Talin^WT^. (**F**-**G**) Confocal images from EC-Talin^WT^ and EC-Talin^KO^ retinas stained for IB4, display the three vascular layers and their colour-coded overlay, (**F**) at P12 following talin deletion on P6,7,8 and (**G**) at P18 following talin deletion on P14, 15, 16. (**H**-**I**) Confocal images of retina whole mount and tumour thick sections from EC-Talin^KO^ mice stained with IB4 lectin to mark blood vessels, display abnormal morphology and balloon-like structures, (**H**) positive for the proliferation marker ki67 and (**I**) negative for the apoptosis marker cleaved caspace 3. Scale bars, 100 μm (**A**-**B**); 20 μm (**C**-**D**, **H**-**I**); 15 μm (**E**); 30 (**F**-**G**); 20 μm. Statistical analysis, Mann–Whitney rank-sum test. Experimental observations of n=20, 6, 10 retinas for **B**, **C**, **E**.

To investigate the talin-dependent steps of the angiogenic process, we exploited the retina angiogenic model where morphogenesis of the retina vasculature occurs in a time-regulated and specific pattern. Initially, the retina vasculature expands from the optic nerve towards the periphery to form the superficial plexus by postnatal day 7 (P7). Following the initial outgrowth, the retinal vessels grow perpendicular to form the deep layer by P12 and subsequently the intermediate layer by P18 (*48*). All three layers besides vascular network expansion, undergo extensive remodelling, including pruning and branching (*49, 50*). To determine the effect of endothelial loss of talin in angiogenic sprouting, we induced talin deletion by injecting 4-OHT in *Pdgfb-*iCreER^T2^; Talin1^fl/fl^ mice on three subsequent days (P1, P2, P3) and examined the retina vasculature at P6. The vascular network in retinas of EC-Talin^KO^ mice was less dense, and the distal end of the vascular plexus appeared blunt (Fig. 2B). In line with a recent report (*51*), loss of endothelial talin severely impaired radial vessel outgrowth (Fig. S4A) and reduced vessel branching (Fig. S4B). The number of endothelial sprouts per angiogenic front was significantly decreased (Fig. S4C). Similar defects in angiogenic sprouting and decreased vessel density and branching were observed in both arterial and venous front (Fig. S4D). Detailed examination of the retina angiogenic front revealed severe vascular malformations that disrupted the architecture of blood vessels. Loss of endothelial talin caused EC aggregation, creating abnormal round cellular arrangements and balloon-like bulging structures (Fig. 2C-D). These morphologically distorted cells were loosely associated with the vascular network, as demonstrated by immunostaining for the endothelial marker Erg and BS1-lectin in P6 and P12 retinas. Shedding of cell clusters or individual deformed cells was evident at the angiogenic front, and the newly-developed vascular networks (Fig. 2C). Tip-stalk cell selection was not affected as no differences were detected in the expression of ESM or Delta-like 4, both established markers for tip cells (Fig. S5A-B) (*52, 53*). Congruent with the presence of tip cells, filopodia were evident in the abnormal vascular structures caused by endothelial talin deletion. However, their length was severely reduced (Fig. 2D), suggesting a role for endothelial talin in regulating actin protrusions. Similarly, tip cells with filopodia extensions were identifiable during the formation of the intermediate layer (Fig. 2E).

To examine the impact of talin deletion on endothelial cell invasion *in vivo*, we induced talin deletion after the formation of superficial plexus by three 4-OHT injections at P6, P7, P8, and examined the retinal vasculature at P12. Detail analysis of the perpendicular sprouting of the superficial vessels revealed blunted ends in retinas from EC-Talin^KO^ and the inability of ECs to invade and form the deep retina layer (Fig. 2E-F) of both arterial and venous origin (Fig. S6). Next, we examined the impact of endothelial talin deletion on the remodelling of the vasculature, following the formation of all layers. We deleted endothelial talin by 4-OHT administration at P14, P15, 16 and examined the retina vasculature at P18. We observed a significant reduction in the vascular area of the deep layer and the newly formed intermediate layer (Fig. 2G) both in arterial and venous vasculature (Fig. S7). Taken together these data demonstrate that endothelial talin is essential for angiogenic sprouting, branching, and remodelling of the vascular network.

The vascular morphological distortions caused by loss of endothelial talin, were detected, also, in tumour blood vessels (Fig. 2H-I). Notably, ECs forming these abnormal round aggregates were viable, as determined by the presence of proliferation marker Ki67 (Fig. 2H) and the lack of active-caspase-3 immunostaining (Fig. 2I). Moreover, these abnormal vascular deformities could be surrounded by NG2-positive pericytes (Fig. S5C). In summary, these findings establish that loss of endothelial talin severely disrupts vascular morphology, causing EC aggregation and balloon-like abnormal structures. These vascular malformations diminished endothelial cell migration and invasion to form angiogenic sprouts, disrupted vessel branching and impaired the remodelling and the stability of newly formed vascular network.

### Loss of talin from well-spread cells, destabilises steady-state adhesion and cytoskeletal dynamics and disrupts cell morphology

To decipher how loss of talin causes vascular deformities and balloon-like EC clusters, we performed detailed characterisation of EC dynamics *in vitro*. Existing studies address the role of talin in cell attachment and spreading using cells where talin is constitutively disrupted (*21, 33, 37*-*41, 54*). This reveals the role of talin during the initial stages of adhesion development but not steady-state adhesion dynamics. Other studies have explored the role of reducing talin protein, using siRNA-depleted cells in two-dimensional substrates (*55, 56*). These approaches, although informative, preclude our understanding of how talin regulates adhesion dynamics *in vivo*, where cells are embedded within tissues. The inducible talin deletion system we use gives us the opportunity to examine the effect of talin deletion following the establishment of cell adhesion in ECs. We isolated primary lung ECs from *Pdgfb-*iCreER^T2^; Talin1^fl/fl^ mice, treated them with 4-OHT (KO) or vehicle control (wt), after cell spreading and the stabilisation of cell adhesions and verified talin’s deletion by western blot (Supplementary figure 8a). To follow the time-course of the effect of talin deletion, we took advantage of the *Pdgfb-*iCreER^T2^; Talin1^fl/fl^; ROSA^mT/mG^ mice that carry the reporter mT/mG cassette in the *ROSA* genomic locus. All wild-type cells in these mice express the mTomato protein and display red fluorescence. Upon 4-OHT administration, the cre-expressing cells switch off mTomato expression and turn on GFP expression, turning their fluorescence from red to green. Using the mT/mG reporter and talin immunostaining we observed remarkable changes in cell morphology upon talin deletion. Cells lost their flat shape and became irregular and spindle-like, as talin was eliminated, finally adopting a round morphology. Endpoint quantification of the cell area revealed that loss of talin caused significant cell shrinkage (Fig. 3A). Similarly, removal of talin from established EC colonies induced cell aggregation and dramatically decreased the colony area (Fig. 3B). Deletion of talin from well-formed endothelial monolayers, caused cell retraction, diminished cell spreading and destroyed cell-cell adhesions (Fig. S8B). Loss of endothelial talin specifically disrupted junctional integrity and decreased expression and localisation of VE-cadherin at cell-cell junctions, as determined by immunostaining and western blot analysis (Fig. S8B-C). To get a clearer insight into the dynamic of cell shape changes elicited by loss of talin, we performed time-lapse microscopy of mT/mG ECs. Talin KO ECs were unable to maintain their cell geometry, unlike wt ECs. Loss of talin caused ECs to undergo rounds of cell stretching followed by abrupt detachment and cell shrinkage. Membrane bulging followed by sporadic re-attachment was also frequently observed (movie S3). In endothelial cell colonies, loss of talin induced cell contraction, decreasing colony area compared to wild-type (movie S4). Likewise, significant cell shape malformations were observed when cells were treated with 4-OHT after spreading in 3-dimensional (3D) collagen gels and stained with phalloidin to visualise F-actin. Loss of talin disrupted the bipolar morphology of endothelial cells, and caused cell rounding, decreasing significantly cell volume (Fig. 3C). Real-time imaging of cell morphology in 3D collagen gels demonstrated that loss of talin distorted EC shape and disrupted membrane protrusion orientation and stability compared to wt cells (movie S5). ECs lacking talin extended long finger-like membrane protrusions that were highly mobile and were retracted abruptly due to their poor anchorage to ECM (movie S6). Congruent with the retina vasculature phenotype, deletion of talin after the formation of vessel-like structures in 3D collagen gels caused cell accumulation and rounding at the distal parts of branches (Fig. 3D).

**Fig. 3.**
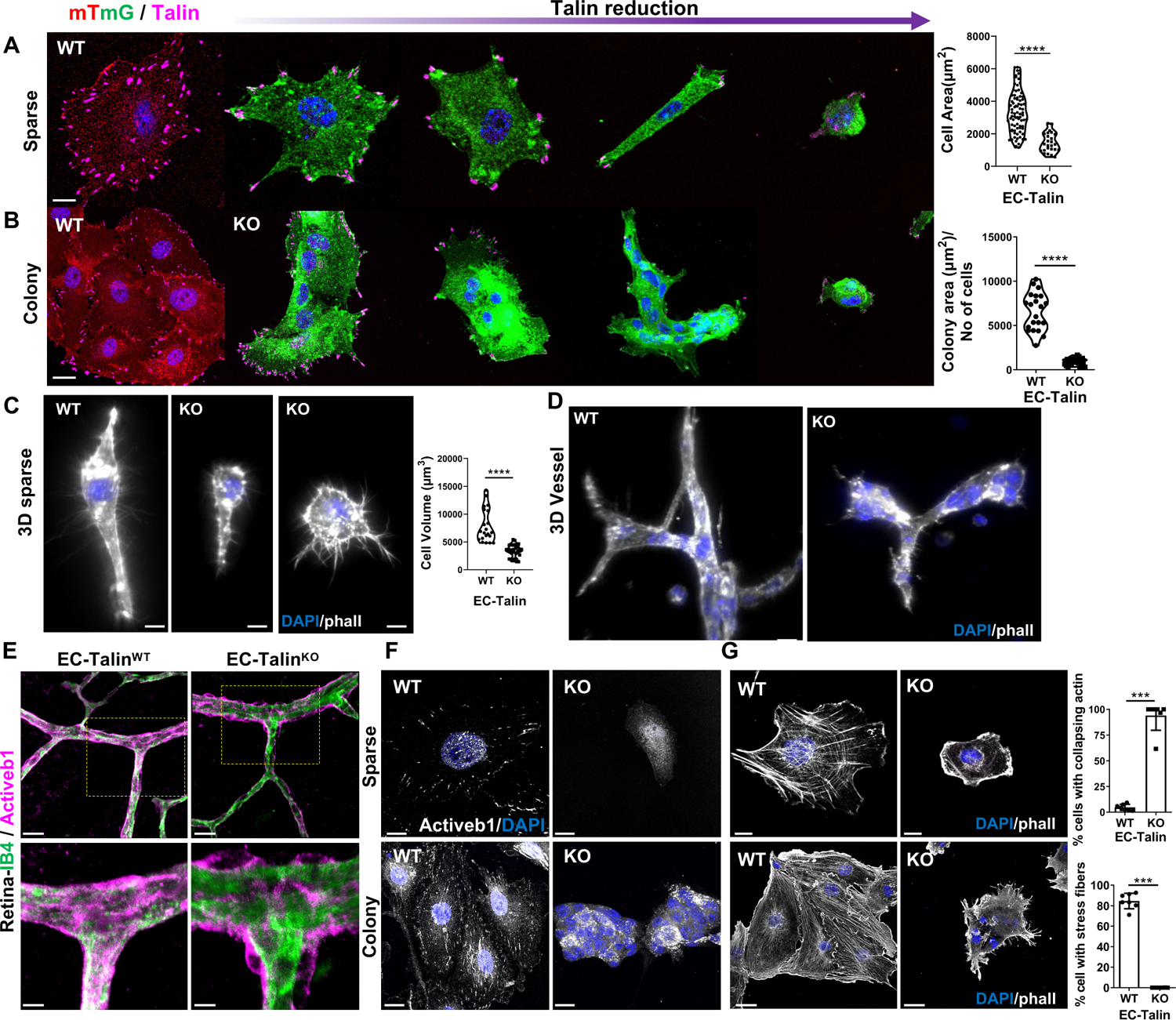
Talin deletion in well-adherent endothelial cells destroyed cell morphology, adhesion and actin cytoskeleton. (**A-B**) Representative fluorescent images of primary ECs isolated from *Pdgfb-*iCreER^T2^; Talin1^fl/fl^; ROSA^mT/mG^ mice and stained with antibody against talin. Disruption of cell shape was observed after the administration of 4-HOT in well-adherent (**A**) sparse cells and (**B**) colonies. Violin plots represent the cell area of individual wt and KO cells and the colony area divided by the number of nuclei in each colony; n=80 wt and 25 KO, for cell area and n=21 wt and 23 KO, for cell colonies, ****P<0.0001. (**C**) Representative light sheet microscopy images of primary ECs embedded in 3D collagen gels and stained with phalloidin showing disrupted cell morphology. 4-OHT induced talin deletion following ECs spreading in 3D. Violin plots represents the cell volume in individual WT and KO cells; n=21 wt and 30 KO, ****P<0.0001. (**D**) Representative light sheet microscopy images of vessel-like structures formed in 3D collagen gels by control (wt) and talin KO ECs stained with phalloidin, showing formation of EC clusters upon deletion of talin after cell spreading. (**E**) Confocal images of EC-Talin^WT^ and EC-Talin^KO^ retinas immunostained for active b1-integrin and isolectin-B4 (IB4) to mark ECs. Insets shows diminished levels of active b1-integrin in ECs and normal levels in surrounding cells; n=10 for each genotype. (**F**) Representative images of primary sparse ECs and colonies showing lack of active b1-integrin staining upon talin deletion from well-adherent cells. (**G**) Representative images of phalloidin-stained sparse ECs and colonies shows disorganization of the actin cytoskeleton following deletion of talin from well-adherent cells (KO). Bar charts represents the percentage of cells with collapsing actin and stress fibers in wt and talin KO ECs, mean ± s.e.m.; n= 7 independet experiments (total cell numbers 380 wt and 183 KO), ***P=0.0006. Nuclei were stained with DAPI in **A-D** and **F-G**. Scale bars, 10 μm (**A, C, F**/sparse**, G**/sparse); 20 μm (**B**, **D**, **E, F**/colony, **G**/colony); 5 μm (insets in **E**). Statistical analysis, Mann–Whitney rank-sum test. Experimental observations in **F** from 20-40 cells and colonies from 3 independent experiments.

Previous studies have demonstrated reduced numbers of cell-matrix adhesions upon talin reduction (*21, 33, 38, 41, 55*). We observed a specific localisation pattern of talin at cell-matrix adhesions. As talin levels were reduced, the remaining talin protein concentrated at a few sparse adhesion sites at the cell periphery, placed distantly to each other, forming the anchor points of the cell (Fig. 3A). Limited cell attachment to only two points, confined talin-labelled adhesions to opposite ends (Fig. 3A). The distinct localisation of cell-matrix adhesions upon talin elimination was also evident in endothelial colonies, where talin labelling was restricted to a few points at the colony periphery, highly reminiscent of the cornerstone adhesions of stem cell colonies (*57*) (Fig. 3B). Immunofluorescent staining for several cell-matrix adhesome members, revealed a co-dependence of talin levels and localisation of vinculin, ILK, pY31-paxillin and FAK at the limited cell-matrix adhesions in sparse cells and in colonies (Fig. S9).

Given the well-established role of talin in integrin activation, we next examined whether removal of talin affected the already engaged integrin at cell-matrix adhesions. Immunostaining of endothelial colonies or sparse ECs following talin deletion by 4-OHT, revealed the complete absence of active b1-integrin detected with the 9EG7 antibody in talin KO compared to wt cells (Fig. 3E). In agreement with a previous study (*44*), we observed reduced activation of b1-integrin *in vivo*. Immunostaining of EC-Talin^KO^ retina vasculature clearly showed that active b1-integrin was present in the surrounding pericytes but completely lost from ECs, labelled by isolectin B4 (Fig. 3F). The loss of b1-integrin activation was not caused by changes in the expression of integrins, because the total levels of b1-integrin were not different between talin-deficient and control cells, as determined by FACs and western blot analysis (Fig. S10A,B). Similarly, no differences in the surface levels of b3-integrin and a5-integrin were observed (Fig. S10B). Outside-in activation also remained unimpaired in talin KO ECs. Thus, manganese (Mn^2+^) treatment, used to induce a high affinity conformation by facilitating unbending of integrins and unclasping of integrin heterodimers (*58*) increased the activation of b1-integrin in talin KO ECs by two-fold, similar to the increase seen in Mn^2+^ treated wt cells (Fig. S10C).

Then we examined whether deletion of talin in well-adherent ECs, affected the cell cytoskeleton. Phalloidin staining of spread ECs and endothelial colonies treated with 4-OHT revealed a complete collapse of the actin cytoskeleton (Fig. 3G). Loss of talin eradicated both actin stress fibres and cortical actin and increased intracellular actin aggregates, without affecting total actin levels (Fig. 3G and Fig. S11A-C). We followed actin dynamics in real-time by transfecting cells with Life-Act GFP and treating them with 4-OHT or vehicle control. Loss of talin did not abolish the ability of cells to form actin fibres but caused a disarray of small membrane protrusions, actin aggregates and disorganised actin cytoskeleton (movie S8) compared to control cells (movie S7). Progressive loss of talin also deregulated actomyosin dynamics. Immunofluorescent staining of phospho-myosin light chain (p-MLC) revealed a highly localised increase in actomyosin contractility at the cell periphery during the initial stages of the talin KO phenotype (Fig. S11B). At the end stages when the actin cytoskeleton was completely disrupted and cells were rounded, p-MLC was undetectable by immunostaining (Fig. S11B). Taken together these findings establish that endothelial loss of talin from stably-attached cells disrupt adhesion and actin cytoskeletal dynamics and destabilise cell shape. This severely impairs the morphology of confluent endothelial monolayers, cell colonies and sparse ECs in two-dimensional and three-dimensional (3D) environments.

### Talin is indispensable for endothelial cell mechanosensing *in vitro* and *in vivo*

Emerging data indicate that talin is an important mechanotransducer (*27, 28*). Our live cell imaging and immunostaining demonstrate the inability of talin KO ECs to stabilise their attachments and to maintain their morphology, pointing to a role of talin in steady-state mechanosensing. To examine whether loss of talin from adherent ECs with established tensional homeostasis, impairs their ability to sense the rigidity of their microenvironment, we seeded primary endothelial cells from *Pdgfb-*iCreER^T2^; Talin1^fl/fl^ mice in fibronectin substrates of different stiffness, ranging from 2-29kPa, and then induced talin deletion with 4-OHT. As expected in soft substrates, both talin-deficient and wt ECs were unable to spread. At 5kPa (*33, 39, 59*), wild-type cells expanded their cell area and formed actin fibers. In contrast, talin-deficient cells adopted a less spread morphology, highly reminiscent of that in soft substrates. Moreover, increasing matrix stiffness did not enhance spreading of talin-deficient ECs (Fig. 4A). These findings directly demonstrate that talin KO cells are unable to sense and respond to different mechanical microenvironments. An established indicator of efficient cell mechanotransduction is the translocation of YAP from the cytoplasm to the nucleus (*60, 61*). To investigate whether loss of talin from cells that are under tensional equilibrium affected mechanotransduction, we induced deletion of talin in well-formed and stable endothelial colonies and sparse ECs. Immunofluorescent analysis of talin-deficient ECs and colonies revealed a wide-spread localisation of YAP, mainly in the cytoplasm with an absence of nuclear accumulation (Fig. 4B). Quantification of YAP immunofluorescent intensity, demonstrated that loss of talin, significantly reduced the nucleus to cytoplasmic ratio of YAP protein (Fig. 4C), without altering total protein levels, as shown by western blot analysis (Fig. 4D). In agreement with this, real-time PCR analysis revealed decreased expression of two known YAP targets, Ankrd1 and Cyr61 (*62*)(Fig. 4E). To validate our findings *in vivo*, we performed immunofluorescent staining of YAP and Erg, an endothelial-specific nuclear transcription factor (*63*), in P6 retinas from EC-Talin^KO^ and control EC-Talin^WT^ mice. Examination of the retina vascular network revealed a variety of mechanosensing stages of wt EC in blood vessels, with some displaying more YAP nuclear staining than others (*64*). EC talin deletion decreased the localisation of YAP in Erg-stained endothelial nuclei, in both the large and small vessels of retina vascular network (Fig. 4F). Collectively, these data established that loss of talin severely disrupts the ability of EC to sense the rigidity of their surroundings in ECs and blood vessels.

**Fig. 4.**
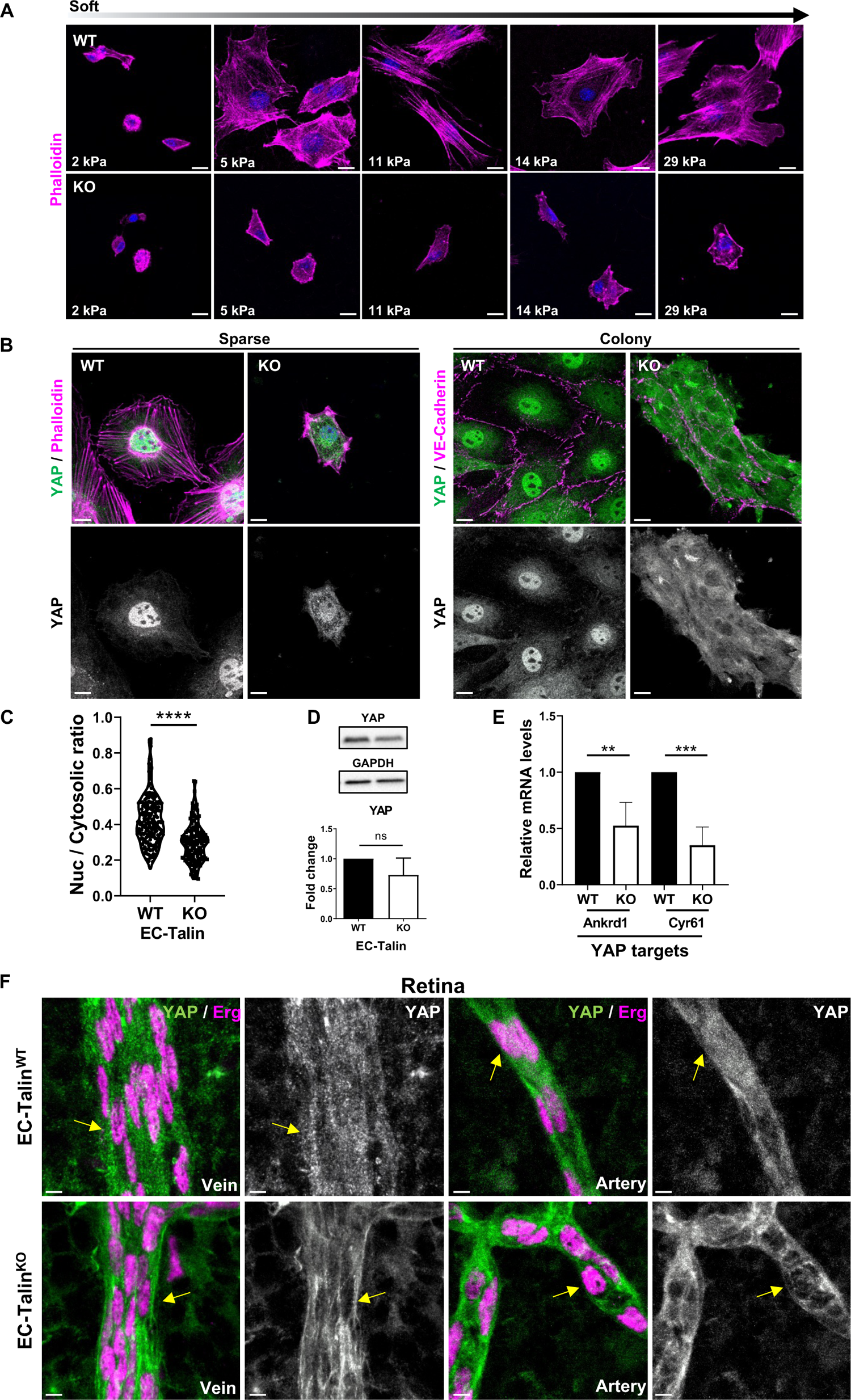
Talin is indispensable for sensing the environmental stiffness in mechanically stabilised ECs and blood vessels *in vivo*. (**A**) Confocal images of F-actin immunodetection by phalloidin in primary control (wt) and talin KO ECs cultured on substrates with different stiffness (2 kPa to 29 kPa). Talin deletion was induced by 4-OHT following cell spreading. Note the reduced cell area of talin KO ECs irrespectively of the matrix rigidity. DAPI marks the nuclei. (**B**) Immunofluorescence images of YAP staining, showing decreased YAP translocation to the nucleus in both talin KO sparse ECs and colonies compared with control (wt). DAPI counterstain nuclei. Phalloidin staining and VE-cadherin immunodetection marked actin cytoskeleton and cell borders, respectively. (**C**) Violin plots represent the ratio of nuclear/cytosolic YAP in wt and KO sparse ECs, n=217 wt and 154 KO ECs from 3 independent experiments, ****P<0.0001, Mann–Whitney rank-sum test. (**D**) Western blot analysis showed no difference of YAP expression levels in wt and KO cell extracts. GAPDH acted as a loading control. Bar chart represents the fold change of YAP protein levels from 3 independent experiments, wt values set to 1, KO: mean ± s.e.m., ns, no statistically significant difference. Two-sided Student’s *t*-test. (**E**) Quantitative PCR analysis revealed significant downregulated mRNA levels of candidate YAP target genes (Ankrd1 & Cyr61) in KO compared to wt ECs. RPLP1 was used as an internal control. Bar chart represents fold changes in mRNA expression (ýýCt) from 3 biological samples (wt values set to 1, KO: mean ± s.e.m.), ***P=0.0002 (Cyr61), **P=0.0037 (Ankrd1). (**F**) Confocal images of veins and arteries from EC-Talin^WT^ and EC-Talin^KO^ retinas immunostained for YAP and Erg to mark specifically endothelial nuclei. Arrows show a uniform YAP localization both in nucleus and cytoplasm of ECs in EC-Talin^WT^ retinas and a reduced nuclear YAP localization in EC Talin^KO^ ECs. Experimental observations from at least n=10 per genotype. Scale bars, 15 μm (**A**); 20 μm (**B**/colony); 10 μm (**B**/sparse, **F**/Arteries); 5 μm (**F**/Veins).

### Expression of the Talin head or Talin rod domains are equally effective in balancing traction forces in endothelial cells

To gain insight into the mechanisms by which loss of talin limits the mechanical responses of ECs and blood vessels, we examined whether different domains of talin protein are sufficient to balance forces and rescue the destabilising effect of talin loss. The ability of talin to bind actin in two different sites in rod domain, has been shown to be necessary for both transmitting traction forces and mediating vinculin mechanotransduction in fibroblasts (*33, 34, 37, 39*-*41*). Furthermore, the talin head domain contains an integrin binding site and has been shown to mediate integrin activation and clustering (*23, 65, 66*). Therefore, we transfected primary endothelial cells from *Pdgfb-*iCreER^T2^; Talin1^fl/fl^ mice with different constructs expressing talin head (1-433aa), talin rod (434-2541aa) and full length talin (1-2541aa) fused to GFP (Fig. S12), and seeded them on fibronectin-coated polyacrylamide gels of approximately 11kPa stiffness, containing fluorescent beads for traction-force experiments. Following treatment with 4-OHT, we analysed bead displacement and traction force levels, using established algorithms (*67, 68*). Loss of talin from well-spread ECs with established adhesions disrupted the tensional equilibrium and traction force generation (Fig. 5A). Expression of the talin rod alone in talin-deficient ECs increased bead displacement and force generation to the level achieved by full-length talin (Fig. 5A). Unexpectedly, we observed that the talin head alone was also able to restore force generation in talin-deficient ECs (Fig. 5A). To corroborate these studies, we performed immunostaining analysis of YAP nuclear translocation in talin-deficient ECs expressing talin-head, talin-rod, and talin-full length. Expression of either the talin-head or talin-rod was able to induce YAP translocation to the nucleus similar to talin-full length expression in talin-deficient cells and to the same extend as wt ECs (Fig. 5B, C). Next, we performed FACs analysis and verified that talin-head was able to increase active b1-integrin levels similar to talin-full length expression whereas talin-rod was unable to rescue b1-integrin activation (Fig. S13). Quantification of cell area demonstrated that expression of either talin-head or talin-rod alone was able to limit the reduction in cell area elicited by loss of talin. However, transfected ECs still occupied less area compared to talin-full length expressing ECs (Fig. 5D). Immunofluorescent examination of actin cytoskeleton by phalloidin staining showed that talin-head and talin-rod alone were sufficient to ameliorate the disorganisation of actin network elicited by loss of talin (Fig. 5E). These data suggest that both talin-head and talin-rod domains are self-sufficient for maintaining cell morphodynamics. Collectively, our findings demonstrate that loss of talin diminishes mechanotransduction in mechanical stable and well-spread ECs. Talin mechanosensing capabilities are linked both to talin head and talin rod domains, suggesting redundant roles of talin head and talin rod in steady-state adhesions. Remarkably, the restoration of traction forces by talin head in talin-deficient ECs provides the first insight into an important and understated role for this domain in mechanotransduction.

**Fig. 5.**
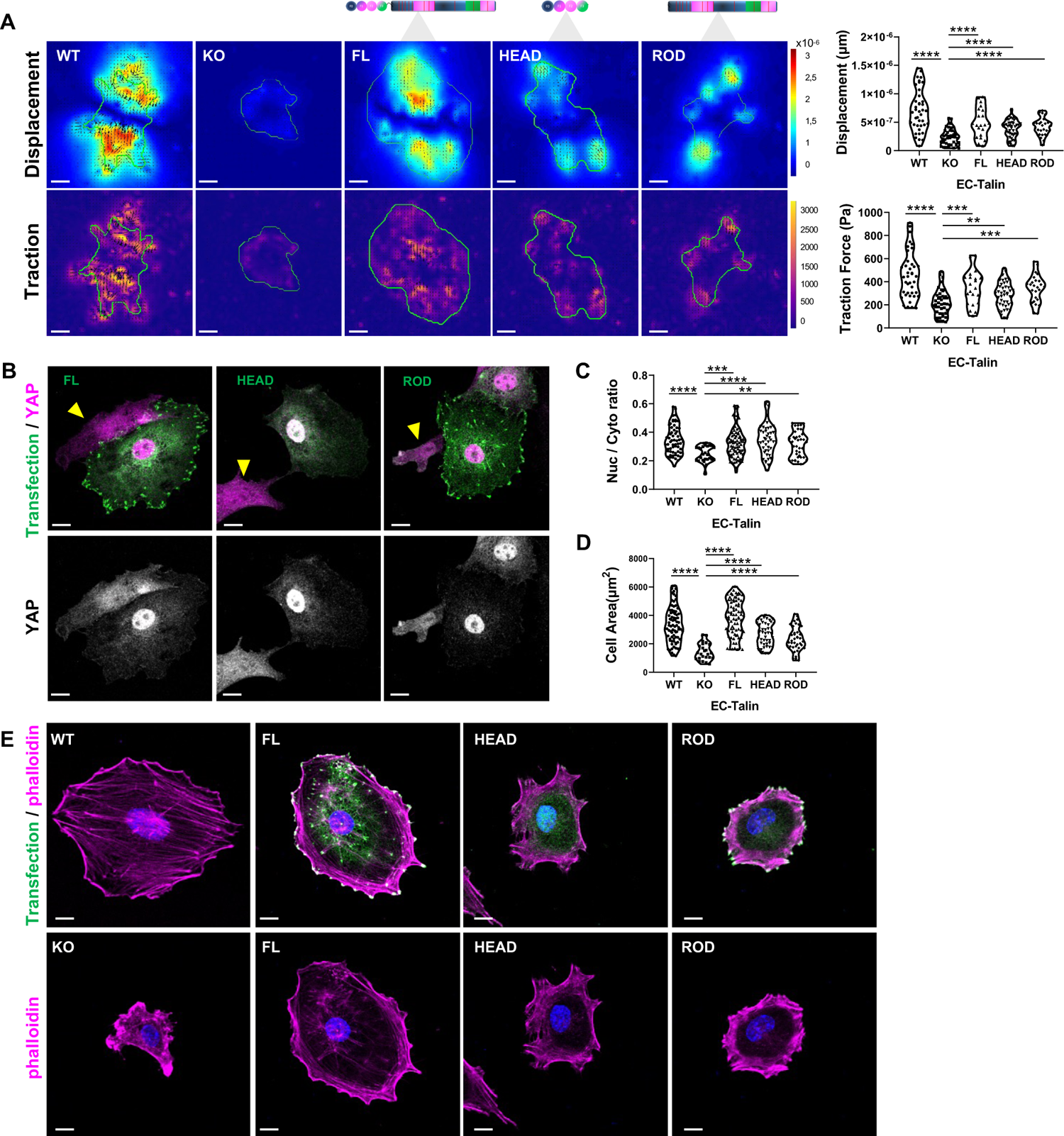
Talin head and talin rod domains play redundant roles in force transmission. (**A**) Representative images of bead displacement and stress magnitude (Pa) from primary control (wt), talin KO ECs and talin KO ECs re-expressing full-length talin (FL), talin head (HEAD) and talin rod (ROD) domains, as indicated in talin-schematic representations. Images are colour-coded on the same scale between the different manipulations, with areas of low displacement in blue and high displacement in red and low traction in purple and high traction in yellow. Note the reduced bead displacement and stress magnitude in talin KO ECs compared to wt controls and the restoration of traction forces upon re-expression of full-length talin, talin head and talin rod domains. Violin plots represents the quantification of mean displacement of beads per cell (upper graph) and mean traction force per cell (bottom graph); n=40, 41, 21, 43 and 25 for wt, KO, FL, HEAD and ROD ECs, respectively. Displacement of beads: ****P<0.0001; Traction force: ****P<0.0001 (WT/KO), ***P=0.0008 (FL/KO), **P=0.0058 (HEAD/KO), ***P=0.0001 (ROD/KO). (**B**) Representative images for YAP immunodetection in talin KO ECs re-expressing full-length talin-GFP, talin-head-GFP and talin-rod-GFP display restoration of YAP translocation to the nucleus. Arrowhead points to untransfected talin KO cells. (**C**) Violin plots represents the nuclear/cytosolic YAP ratio in wt, talin KO, and talin-KO ECs re-expressing full-length talin (FL), talin head (HEAD) and talin rod (ROD) domains. n=60, 26, 78, 44 and 37 for wt, KO, FL, HEAD and ROD ECs, respectively. ****P<0.0001 (wt/KO), ***P=0.0005 (FL/KO), ****P<0.0001 (HEAD/KO), **P=0.0040 (ROD/KO). (**D**) Violin plots represents the cell area of wt, talin KO, and talin-KO ECs re-expressing full-length talin (FL), talin head (HEAD) and talin rod (ROD) domains. n=80, 25, 75, 42 and 38 for wt, KO, FL, HEAD and ROD ECs, respectively. ****P<0.0001. (**E**) Confocal images of F-actin immunodetection by phalloidin staining corresponding to wt, KO, and cells expressing Talin-FL, Talin-HEAD and Talin-ROD domain respectively. DAPI counterstained the nuclei. Scale bars, 10 μm. All experimental observations are from at least 3 independent experiments. Statistical analysis, Mann–Whitney rank-sum test.

## Discussion

Our findings provide *in vivo* and *in vitro* evidence by which talin controls endothelial mechanical stability to drive and sustain the growth of vascular network and fuel cancer progression. To our knowledge, this is the first study that links the mechanical function of talin to a pathological outcome in a mammalian organism. Specifically, we show that deletion of talin abolishes the ability of ECs to sense tissue rigidity and balance cellular forces, leading to disruption of EC shape and connectivity. The resulting vascular malformations and malfunctions *in vivo*, not only blocked the initial stages of tumour development, but more importantly for the clinic, caused regression of established tumours.

The *in vivo* prerequisite of talin mechanosensing to control blood vessel morphology and sustain cancer development, was further supported by the reduced nuclear translocation of the established mechano-effector YAP (*60*-*62*), following endothelial talin deletion. Endothelial YAP/TAZ has been shown to regulate angiogenesis and drive tumour progression (*69, 70*). In a recent study, mechanical cues were found to induce YAP/TAZ nuclear localisation in ECs and increase metastatic angiogenesis, in the absence of chemical signals (*71*). The molecular mechanism involved in rigidity-triggered YAP/TAZ activation is not fully-elucidated. FAK/Src signalling, Rho-GTPases and actomyosin contractility are reported to mediate YAP activation (*62, 70*). Our findings suggest that talin constitutes a central player of the molecular machinery by which YAP/TAZ sense and is regulated by stiff matrices, acting as an important sensor that translates microenvironmental mechanical cues to cellular responses. This outcome has great potential for manipulating YAP activity in cancer treatment.

The tumour vasculature is both affected and contributes to mechanical properties of tumour microenvironment (*72*). ECs are key players in sensing and responding to hemodynamic forces physiologically generated by blood flow and tensional strain created by pathological intratumoural pressure (*17, 73*). Much of our knowledge about EC mechanoresponses during vascular development and homeostasis stems from experiments on adherens junctions (*74*-*76*). Our data show that cell-matrix stiffness can also induce global mechanoresponses and suggest that talin is a central regulator of the mechanical cross-talk between cell-cell and cell-matrix adhesions. Recent studies show that talin, similar to b1-integrin, parvin and vinculin, is required to stabilise VE-cadherin at endothelial adherens junctions (*44, 77*-*79*). Consistent with this, endothelial talin deletion in established monolayers, disrupted cell-cell junctions and increased permeability in tumour vasculature, *in vivo*. Strikingly, the vascular deformities induced by endothelial loss of talin were more severe than previously reported by deletion of other adhesome components (*15*), including ILK (*80*-*82*), FAK (*83*-*85*) and parvin (*79, 86*), suggesting that talin is a major regulator of blood vessels. Moreover, our studies revealed striking vascular deformities with balloon-like EC clusters in both veins and arteries, in contrast to observations on endothelial b1-integrin deletion (*78, 87*). Given the overlapping functions of integrins (*88, 89*) and the important role of b3-integrin in tumour angiogenesis (*90*), our data indicate that talin acts as an integrator of integrin function to orchestrate blood vessel development and function.

Talin has an established role in the formation and reinforcement of the nascent adhesions during cell spreading (*27, 28*), whereas its function in mature adhesions is poorly understood. Previous studies have found that talin cleavage by calpain can regulate adhesion disassembly (*91, 92*). Moreover, talin has a slow turnover rate at established adhesions (*93*-*95*), suggesting a role in maintaining the molecular structure of cell-matrix adhesions. Our findings provide the first direct evidence for the requirement of talin mechanosensing to preserve the molecular architecture of cell-matrix adhesions. Deletion of talin in well-spread and stably-attached ECs, dismantle their adhesion sites, forming anchor-like adhesions reminiscent with the cornerstone adhesions of stem cells (*57*). Additionally, our live imaging data, revealed, that although actin filaments are formed in the absence of talin, talin is necessary for preserving cell cytoskeletal organisation. Consistent with this, filopodia were present at retina tip cells *in vivo*, and albeit shorter in length, were increased in number upon talin deletion. Explanations for the observed collapsing cytoskeleton triggered by talin elimination include destabilisation of adhesion composition or disruption of talin’s direct binding to actin fibers. Collectively, our findings suggest that talin is indispensable for steady-state adhesion and cytoskeletal dynamics.

Much of our current understanding of how talin senses the rigidity of substrates and affects cell attachment, comes from experiments using cells with constitutive talin deletion, which limits studies to the initial events of cell spreading and formation of early adhesions (*21, 33, 34, 37*-*41, 54, 56, 95, 96*). The novelty of our findings stems from examining the effect of talin deletion in steady-state mechanically balanced cells which are well-spread with established adhesions, resembling more the *in vivo* condition. Such studies are more relevant to how cells experience and respond dynamically to forces during homeostasis, tissue regeneration and pathological situations of mammalian organisms. From a clinical perspective, gaining molecular insight into mechanotransduction *in vivo*, could pave the way for the development of novel therapeutic interventions. Despite the vital importance of mechanical forces in all cellular functions, the molecular mechanisms regulating dynamic mechanoresponses in tensional equilibrated environments remain unresolved (*97*). Besides pharmacological inhibition of integrin-mediated signalling (*95*) and actin cytoskeleton (*93, 94, 98, 99*), to our knowledge, this is the first study using specific gene targeting to eliminate a key integrin mechanosensor and establish its role on preserving rigidity-triggered responses. Our findings revealed that talin constitutes an irreplaceable mechanosensor, required for securing steady-state tensional homeostasis and dynamically controlling cellular responses to forces, which safeguard cell shape, adhesion, motility and survival.

A major outcome of our studies is the discovery of the redundant role for talin head and talin rod in mediating talin mechanosensing in stably attached ECs. Cellular studies using constitutively deleted talin fibroblasts found that neither talin head nor talin rod could rescue the defective force transduction in talin depleted fibroblasts during initial cell spreading and adhesion growth (*39*), albeit being reported to generate weak forces (*21, 54*). This is in stark contrast to our findings whereas both talin head and talin rod alone were sufficient to transmit traction forces upon elimination of talin in previously well-adherent ECs. This could be explained by diverse molecular requirements for rigidity sensing between well-attached versus initially spreading cells. The geometry of force transmission could be different during the development of adhesion sites and in tensional equilibrated adhesions. According to this, not all talin molecules are under the same tension in the adhesion sites (*24, 33, 34, 100*), suggesting the involvement of other adhesome members, including FAK, paxillin, vinculin, ILK/parvin, zyxin, a-actinin in assisting talin during co-ordination of cell mechanoresponses (*101*-*110*). Our finding that both talin head and talin rod can partially rescue rigidity-triggered responses suggests that talin utilises different scaffolds and signalling molecules to mediate mechanotransduction, adhesion stabilisation and signalling. Consistent with this notion, elegant studies in *Drosophila* have demonstrated a tissue-dependent orientation of talin and revealed the requirement of different talin interactions for mediating integrin adhesion in wing epithelium and in muscle attachment (*111*).

An important question is how talin head and talin rod alone form functional mechanical linkages to integrins and actomyosin contractile machinery to elicit mechanoresponses in ECs. Given the established function of talin in integrin activation and clustering (*23, 65, 66, 112, 113*) and in line with the ability of talin head to cluster integrins independently of a functional actin cytoskeleton (*114, 115*), we observed increased integrin activation upon talin head re-expression in talin-deficient ECs. Thus, we propose that talin head stabilises integrin clusters, facilitating the rebinding of dissociated integrins before leaving the adhesion sites and possible induce further integrin activation. The linkage of talin-head dependent integrin clusters to actomyosin machinery could be mediated by the putative actin binding site that has been reported to exist in talin head (*116*-*118*) with unknown yet function. We can speculate that talin head uses this actin-binding site to re-enforce the association of integrins with the actin cytoskeleton and create a minimal mechanical linkage that can contribute to cellular tension integrity. An alternative link to actin cytoskeleton could form by other actin-binding adhesome components previously located at adhesion sites. Recently, vinculin was found to interact with paxillin in the absence of talin, suggesting an alternative link to actin cytoskeletal, via vinculin-actin interactions (*42*). Furthermore, the organization of actin around talin head-stabilised adhesions by interactions with the formin FHOD1 (*54*) could facilitate force transmission by talin head. Although further studies are needed to address this possibility and putative differences in the requirements for force generation during spreading and in tensional equilibrated cells, our findings indicate an important and context-dependent role of the talin head in mechanotransduction.

Recently, a point mutation in the integrin binding site of talin’s head domain was reported to decrease B16 melanoma growth and angiogenesis (*51*) but did not eliminate tumour growth as we observed upon completely deletion of endothelial talin, indicating a putative compensatory function mediated by talin’s rod action, *in vivo*. It is well established that upon integrin engagement, the talin rod serves as the main mechanosensing module to elicit cellular responses to forces (*28*). Several structural and cellular studies have dissected the contribution of talin’s rod subdomains and their interactions with actin, vinculin, RIAM and DLC in force transmission (*32, 33, 37, 40, 41, 119, 120*). All these studies have been conducted in the presence of talin head to provide connection to integrins. Our findings indicate that the talin rod can sense and transmit forces in talin-deficient cells independently of the talin-head. A plausible linkage to integrin in the absence of talin head, could be provided by the second integrin binding (IBS2) site that is located at the R11-R12 subdomains before the C-terminal actin binding site (*121, 122*). The IBS2 has been reported to interact with activated b3-integrin subunit (*123, 124*). Although the significance of the IBS2 is not well defined, FRET-studies in *Drosophila* have shown proximity to b integrin subunit, suggesting possible roles in wing epithelium (*111*). Our data provide additional evidence for an important role of IBS2 in mediating an alternative connection of talin to integrins. In the context of previously well-spread and stably adherent cells, talin’s rod direct interaction with activated b-integrin subunit and connections with actomyosin machinery create a mechanical linkage sufficient to probe tissue stiffness and exert cellular traction forces. Additional strengthening of integrin adhesions could be mediated by vinculin mechanical function. In line with this, the C-terminal region of talin can recruit vinculin to adhesion sites and activate FAK signalling (*56*). Moreover, recent studies showed that talin-rod subdomain R7 binds directly to the KANK protein family and interacts with microtubule cytoskeleton (*125, 126*) and this connection is regulated by forces (*127*). Such interactions between the talin rod and KANK proteins could regulate force transmission at cell-matrix adhesions and support the mechanical stability of talin-deficient cells. Taken together, in a physiological situation, one could envisage that talin head is involved in bearing forces applied from ECM to integrins and the talin rod is the main contributor engaged in forces produced by actomyosin contractility. Uncoupling these putative two mechanical properties of talin *in vivo* could be the focus of future studies.

Collectively, our findings establish that talin is required for sensing the rigidity of extracellular matrix and transmits forces that control cell morphodynamics in well-attached cells, thus illuminating the undetermined molecular mechanisms that regulate steady-state cellular tension. Importantly, this study reveals for the first time the *in vivo* mechanosensing requirement of talin, describing a vital role in blood vessels and cancer. Additionally, our data challenge the current thinking by which the main mechanosensing module resides in talin rod domain and uncovers an under-appreciated role for the talin head in force transmission and mechanical stabilisation of ECs and blood vessels *in vivo*. The outcome of this study provides the first evidence where loss of a key adhesome member affected established tumours and disturbed vascular function by disrupting the mechanical properties of endothelial cells. Given the importance of tumour mechanics in cancer progression, our findings identify a new possibility of endothelial mechanomodulation that could be exploited in cancer therapeutic strategies. One could envisage manipulating talin’s mechanosensing capability to disrupt the tumour vasculature and diminish established tumours.

## Materials and Methods

### Mice and Ethical regulations

The floxed Talin (Talin1^fl/fl^) transgenic mice (*46, 47*) were bred with *Pdgfb-*iCreER^T2^ transgene (*45*)to generate *Pdgfb-*iCreER^T2^; Talin1^fl/fl^ mice. Inducible endothelial-specific deletion of talin was achieved by oral gavage delivery of three daily doses of tamoxifen, first dose of 4mg/mice and two consequent doses of 2mg/mice. Also, mice were crossed with ROSA^mT/mG^ double-fluorescent reporter mouse (Jackson Laboratory no007676) to visualize cre activity. For retinal angiogenesis experiments, newborn offspring from *Pdgfb-*iCreER^T2^; Talin1^fl/fl^ (males) x Talin1^fl/fl^ (females) intercross received three consequent daily intraperitoneal injections of 4-hydroxy-tamoxifen (4-OHT, purity ≥98% Z isomer, Sigma-Aldrich Cat# H7904) (80 μg/pup diluted in ethanol/peanut oil) on P1-3, P6-8 and P14-16, and retinas were analysed at postnatal stages P6, P12 and P18, respectively. All mice were bred in the animal facilities of BSRC “Alexander Fleming” under specific pathogen-free conditions. All animal procedures and experimental protocols were approved by the Veterinary Administration Bureau, Prefecture of Athens, Greece under compliance to the national law and the EU Directive 63/2010 and performed in accordance with the guidance of the Institutional Animal Care and Use Committee of BSRC Al. Fleming.

### Syngeneic mouse tumour models

Age (8-12 weeks old) and sex matched *Pdgfb-*iCreER^T2^; Talin1^fl/fl^ mice received subcutaneous injections of 1×10^6^ B16F0 melanoma cells, αnd 1×10^6^ CMT19T lung carcinoma cells (ATCC) at the flank. Endothelial talin deletion was achieved at different points of tumour growth by oral gavage administration of three tamoxifen doses on day 0,1,2 for initial growth studies, day 5,6,7 for subsequent tumour growth and day 10,11,12 for established tumour studies and tumours were excised at day 14 for B16F0 and day 16 for CMT19Ts. For thick tumour section analysis, mice were heart perfused with 4% PFA before tumour excision. For tumour regression experiments, tumour dimensions were measured every two days using a caliper. Tumour volume was determined using the ellipsoid formula length × width^2^ × 0.52.

### MicroPET/CT imaging

Whole body microPET/CT scans were performed using Mediso nanoScan PET/CT scanner at microPET/CT unit of Biomedical Research Foundation, Academy of Athens. Briefly, after 15 hours of fasting, each mouse bearing the B16F0 tumour, received intravenously 100ml of 3.7-7 MBq of 2-^18^fluoro-2-deoxy-D-glucose (^18^FDG). During the uptake phase, animals were anesthetized using isoflurane. The CT scan and a 15-min whole body PET scan were performed successively 60 minutes post injection, while anesthesia was maintained with 2% isoflurane in oxygen (2.4l/min). During the uptake phase and acquisition the body temperature of the animals was maintained at 36°C on the scanner’s bed. For CT scan, the energy of X-ray beam was set at 50 kVp, the current at 670 mΑ and the exposure time at 300 ms. The CT image reconstruction was performed using Nucline software version 2.01. The PET data were reconstructed using a version of 3D OSEM algorithm (Tera-Tomo 3D PET image reconstruction) with four subsets and thirteen iterations. The voxel dimensions of PET and CT were 0.4 x 0.4 x 0.4 mm^3^ and 0.25 x 0.25 x 0.25 mm^3^, respectively. All the required corrections (dead time, decay, attenuation and scatter) were applied, and normalization was performed on the PET data. Images were analysed with the software InterView Fusion 3.00.039.0000, manufactured by Mediso. The mice were imaged 8, 10 and 14 days after tumour cell injection. Oral gavage administration of 4mg/mouse tamoxifen was performed on day 10 after the second PET/CT scan, and two doses of 2mg/mouse on day 11 and 12. Last microPET/CT was performed on day 14 of tumour growth.

### Immunofluorescent analysis of tumours

Tumours were snap-frozen or fixed overnight with 4% PFA for 24 h, cryopreserved using 30% sucrose and frozen in OCT. Midline thin cryosections (10 mm) from snap-frozen tumours were fixed in cold acetone for 10 min at −20 °C and for PFA-fixed tumours were post-fixed with cold 4% PFA for 10 minutes and permeabilized with PBS 0.3% Triton X-100 for 10 min. All sections were blocked with 1% BSA in PBS for one hour and incubated with primary antibodies overnight at 4°C. Sections were then washed in PBS and incubated with Alexa Fluor-labelled antibodies (dilution 1:400) for one hour at room temperature. Whole thin tumours sections were imaged using the ZEISS AxioObserver Z.1 microscope and the tile module of ZEN3.2 blue software. Image analysis of antibody-stained area over the total tumour section area was performed using the ImageJ software. For 3D vessel analysis, midline thick free-floating tumour cryosections (50 mm) were stained as above, with the exception of increased permeabilization time to 30 min and overnight incubation of secondary antibodies. Imaging of thick sections was performed with the LEICA Confocal SP8X (WLL) microscope and analysed with IMARIS software. For the analysis of vessel coverage with a-smooth muscle actin-positive cells, 3D surfaces of each staining were created in IMARIS 8.2.1 software. The area of PECAM surface that overlaps with the area of α-smooth muscle surface divided by the total area of PECAM surface was calculated using the surface-surface contact area extension of the software.

To examine the tumour vessel leakage, mice were intravenously injected with 100ul (1mg/ml) 70KDa-FITC-conjugated Dextran (Sigma-Aldrich) 10 minutes before sacrifice. Tumours were snap-frozen and thick (50 mm) cryosections were stained with PECAM (1:400 dilution) to define the tumour vessel boundaries. 3D surfaces representing the PECAM and the FITC-Dextran staining were created using the IMARIS 8.2.1 software and a colocalisation surface of the two stainings was generated using the surface-surface colocalization extension of the software. To define the FITC-Dextran area that was external to the tumour vessels and represented the vessel leakiness, the colocalization surface of PECAM with FITC-Dextran was subtracted from the total FITC-Dextran area and divided with the total FITC-Dextran area.

Tumour hypoxia was detected using HypoxyprobeTM-1 Kit ((Hypoxyprobe™-1 HPI, Inc) following the manufacturer instructions. Tumour-bearing mice received intraperitoneal injection of pimonidazole (60 mg/kg body weight) one hour before culling by cervical dislocation. Tumours were harvested, snap-frozen and cryosectioned. Thin tumour sections (10 μm) were fixed for 10 min in acetone at −20 °C, and incubated with FITC-conjugated anti-pimonidazole antibody (1:50 dilution) overnight at 4 °C. Sections were then washed with PBS and mounted using Mowiol. Fluorescent images were acquired using ZEISS AxioObserver Z.1 microscope. The ratio of hypoxic-stained area over the total area of the tumour section was determined using ImageJ software.

The following antibodies were used: Ki67 (1:300 dilution, Rabbit polyclonal; Abcam, Cat# ab66155), Cleaved Caspase-3 (1:300 dilution, Rabbit monoclonal; Cell Signaling, Cat# 9664), PECAM (1:400 dilution, Rat monoclonal, clone MEC 13.3; BD Biosciences, Cat# 553370), anti-actin, α-Smooth Muscle - Cy3™ (1:500 dilution, Mouse monoclonal, clone 1A4; Sigma-Aldrich, Cat# C6198), F4/80 (1:100 dilution, clone Cl:A3-1; Serotec, Cat# MCA497) and CD11b (1:100 dilution, clone M1/70; BD Biosciences, Cat# 550282).

### Aortic ring assay

Aortic ring assay was performed as described previously (*128*). Briefly, thoracic aortae were dissected from *Pdgfb-*iCreER^T2^; Talin1^fl/fl^ mice, cleaned from surrounding tissues and cut into 1mm rings. Rings were serum starved overnight at 37°C before being embedded in 1.5mg/ml collagen gels and cultured in Opti-MEM™ reduced serum medium (supplemented with 2.5% (vol/vol) FBS and 30 ng/ml VEGF. Rings were fed every other day with fresh medium until the end of the experiment. Talin deletion was achieved by two daily doses of 1mM 4-OHT (Sigma-Aldrich Cat#H7904) post-embedded in day 0,1 for initial growth and day 3,4 for established sprouts. Rings were fixed after 7 days with 2% PFA and stained with rat PECAM (dilution 1:100, Rat monoclonal, clone MEC 13.3; BD Biosciences, Cat# 557355). Rings were imaged in 3D using the ZEISS Lightsheet Z.1 microscope. The number of sprouts were counted under a microscope while the branching points per ring quantified via IMARIS software.

### Whole-Mount Immunostaining of Retinas

Whole animal eyes were collected from littermate pups and were fixed with 4% PFA overnight at 4°C or retinas were freshly dissected following a brief fixation with 4% PFA for 10 min. Dissected retinas were either fixed with 1:1 MeOH/PBS for 1 hour at 4°C and stored in MeOH at −20 C or stained immediately. Retinas were permeabilized and blocked in 1% bovine serum albumin (BSA) with 0.3% Triton X100 overnight at 4 °C. Following washes three times in PBLEC buffer (1 mM CaCl2, 1 mM MgCl2, 0.1 mM MnCl2 and 1% Triton X-100, in PBS) for 20 min, retinas were incubated with biotinylated isolectin B4 (1:50 dilution, biotinylated; Vector Labs, Cat# B-1205), diluted in PBLEC, overnight at 4°C. Retinas were then washed five times with 1:1 blocking buffer in PBS for 20 min and incubated with Alexa Fluor streptavidin-conjugated antibodies (1:100; Life Technologies) for 2 hours at room temperature. Following four washes, retinas were flat-mounted in microscope glass slides with mowiol mounting medium. For double whole-mount immunofluorescent staining, retinas were further incubated with primary antibodies in blocking buffer overnight at 4°C. Following extensive washes in PBS, retinas were incubated with Alexa Fluor-labelled secondary antibodies (1:500) for 2 hours at room temperature and flat-mounted with mowiol mounting medium. Imaging was performed with the LEICA Confocal SP8X (WLL) microscope. The primary antibodies used were: Erg (1:600 dilution, Rabbit monoclonal, clone EPR3864; Abcam, Cat# ab92513), Ki67 (1:250 dilution, Rabbit polyclonal; Abcam, Cat# ab66155), cleaved Caspase-3 (1:100 dilution, Rabbit monoclonal; Cell Signaling, Cat# 9664), YAP (1:100 dilution, Mouse monoclonal; Santa Cruz, Cat# sc-101199), active Integrin b1 (1:100 dilution, Rat monoclonal, clone 9EG7; BD Biosciences, Cat# 550531), ESM-1 (1:100 dilution, Goat polyclonal; Novus, Cat# AF-1810), DLL4 (1:25 dilution, Goat polyclonal; Company, Novus, Cat#AF1389) and NG2 (1:200 dilution, Rabbit polyclonal; Sigma-Aldrich, Cat# AB5320). Quantification of radial outgrowth, branching points and sprouts per angiogenic front was performed as described (*129*) using the ImageJ software. Analysis of the retina vascular layers and quantification of filopodia length was performed using the IMARIS 8.2.1 software. The isolectin B4 staining was used to create and present in single colour code the surfaces of each vascular network layer. The total surface area of the vasculature in each retina layer was measured.

### Primary 2D and 3D endothelial cell culture

Primary mouse lung endothelial cells (MLECs) were isolated from *Pdgfb-*iCreER^T2^; Talin1^fl/fl^ mice and cultured as described previously (*130*). MLECs were plated in pre-coated tissue culture plates with a mixture of collagen (30mg/ml), gelatine (0.1% in dH_2_0), and fibronectin (10mg/ml) and allowed to attach and spread at least overnight. Talin deletion occurred by administration of 5mM 4-OHT (purity ≥98% Z isomer, Sigma-Aldrich Cat#H7904) post-spreading. For 2D live cell imaging, cells were plated onto glass-bottom 4-well m-Slides (ibidi) pre-coated with collagen and fibronectin mix and allowed to spread. Live imaging was started 60 hours after the initial addition of 4-OHT and maintained for 24hours. In this time window we were able to observe the development of the phenotype induced by talin deletion. Images were acquired every 10 min with a 40X lens using ZEISS AxioObserver Z.1 microscope. For 3D cell cultures, cells were embedded in 1mg/ml rat tail derived type I collagen (Corning) and casted in homemade agarose capillaries. 3D collagen cultures, were then treated with 5 mM 4-OHT (Sigma-Aldrich, Cat#H7904) Sigma) to achieve talin deletion. Time-lapse 3D imaging was performed with a 20X WPlan-APO lens every 10 min using ZEISS Lightsheet Z.1 microscope.

### Immunofluorescent analysis of cultured cells

Cell were seeded on 13 mm round glass coverslips pre-coated with collagen and fibronectin mix and allowed to grow either as sparse cells or in cell colonies. Following talin deletion, cells were fixed for 10 min with 4% PFA or for 3 min with ice-cold MeOH and washed with PBS three times. PFA-fixed cells were permeabilized with 0.1% PBS-Triton X-100 for 10 min, washed with PBS and blocked with 1% BSA/PBS for one hour at room temperature. Primary antibody incubations were performed in 0.1% BSA/PBS, overnight at 4°C and secondary antibody incubations in 1% BSA/PBS for one hour at room temperature. Phalloidin was added with the secondary antibodies to stain F-actin and DAPI was used for nucleus visualization. Mowiol mounting medium was used to mount coverslips. Fluorescence imaging was performed using the LEICA Confocal SP8X (WLL) microscope. Cell area in sparse or in colony cultures was calculated with the ImageJ software by drawing the outline of the cells or colonies. For the analysis of the actin cytoskeleton, the number of cells with collapsing actin, stress fibers and cortical actin, as defined in supplementary figure 11, were manually counted and divided by the total number of cells.

The nuclear/ cytosolic ratio of YAP was quantified using the ImageJ software. The nuclear YAP staining was calculated by measuring the signal intensity of YAP staining in nuclei masks created using the DAPI signal intensity. A freehand selection demarcating each cell area was used to measure the total intensity of YAP staining. The cytosolic YAP staining was calculated by subtracting the nuclear YAP from the total YAP staining in each cell.

At the endpoint 3D cell cultures were fixed with 4% PFA for 10 minutes, permeabilized for 30 minutes with 0.5% Triton X-100 in PBS and blocked with 1% BSA in PBS for 1hour. To stain for actin, samples were incubated with rhodamine-conjugated phalloidin (1:400) overnight followed by washes with PBS. DAPI was used for nuclei visualization. Imaging was performed with a ZEISS Lightsheet Z.1 microscope.

The following primary antibodies were used: Active b1-Integrin (1:100 dilution, clone 9EG7; BD Biosciences, Cat# 550531), actin (1:400 dilution, Sigma-Aldrich, Cat# A2103), YAP (1:100 dilution, Santa Cruz, Cat# sc-101199), talin-1 (1:300 dilution, Clone 97H6; Serotec, Cat# MCA4770), talin-1 (1:100 dilution, Cell Signaling, Cat# 4021), vinculin (1:100 dilution, clone hVIN-1; Sigma-Aldrich, Cat# V9264), ILK (1:500 dilution, Cell Signaling, Cat# 3862), phospho-Paxillin (Tyr118) (1:100 dilution, Cell Signaling, Cat# 2541), phospho-FAK (Tyr397) (1:100 dilution, Thermo Fisher Scientific, Cat# 44-624G), FAK (1:100 dilution, clone 4.47; Millipore, Cat# 05-537), kindlin-2 (1:200 dilution, clone 3A3; Millipore, Cat# MAB2617) paxillin (1:100 dilution, clone 349; BD Biosciences, Cat# 610052) phospho-myosin light chain (pMLC) (1:100 dilution, Cell Signaling, Cat# 3671), VE-cadherin (1:100 dilution, BD Biosciences, Cat# 555289).

### Constructs and transfections

All primary endothelial cell transfections were performed by electroporation (NEPA™ transfection system) using 1×10^6^ cells resuspended in 100ul Opti-MEM™ reduced serum medium containing 10 mg plasmid DNA. pLifeAct-mTurquoise2 was a gift from Dorus Gadella (Addgene plasmid # 36201). The full-length pEGFP-talin-1, pEGFP-talin-head (1-433aa) and pEGFP-talin-rod (434-2541aa) plasmids were kindly provided by Prof. David Critchley. The electroporation parameters were set as follows, poring pulse: voltage 150V, length 7.5ms, interval 50ms, number 2, decay rate 10%, polarity ‘+’; transfer pulse: voltage 20V, length 50ms, interval 50ms, number 5, decay rate 40%, polarity ‘+/-’.

### FACs analysis

At the end-point of talin deletion, adherent cells were detached from the plate and stained with primary antibodies diluted in FACs buffer (PBS supplemented with 1 mM CaCl_2_, 1 mM MgCl_2_, 1% BSA) for 30 minutes on ice, washed twice with cold FACs buffer and then incubated with Alexa Fluor-labelled secondary antibodies (dilution 1:400) on ice for 30 minutes. For Mn^2+^ experiments, cells in suspension, were incubated with 5mM MnCl_2_ in Opti-MEM™ reduced serum medium for 10min at 37°C prior to staining. Flow cytometry was carried out with a FACS CantoTM II cytometer (BD Biosciences) equipped with FACS DiVa software (BD Biosciences). Data analysis was performed with the FlowJo program (version 9.4.10). The following primary antibodies were used: active b1-integrin (1:50 dilution, clone 9EG7; BD Biosciences, Cat# 550531), b1-integrin (1:50 dilution, clone MB1.2; Millipore, #MAB1997), b3-integrin-PE conjugated (1:100 dilution, clone 2C9.G2, BD Biosciences, Cat# 553347) and a5-integrin (1:50 dilution, clone BMC5; Millipore, # MAB2575).

### Preparation of polyacrylamide hydrogel with gradient in stiffness

Polyacrylamide gels with stiffness gradients between 2kPa and 29kPa were prepared according to the protocol adapted from (*39*). Different concentrations of acrylamide and bis-acrylamide were mixed in a solution containing 0.5% ammonium persulfate (Merc Millipore), 0.1% tetramethyl ethylenediamine, TEMED (Fisher Scientific). Ten ml of each mix was placed on a silanized with 0.5% 3-aminopropyltrimethoxy-silane (ACROS) and activated with 0.5% glutaraldehyde (Sigma) glass-15mm coverslip and covered with a 13mm coverslip pre-coated with fibronectin (1mg/ml). Gels were allowed to polymerize for 30min, then the top coverslip was removed and the polyacrylamide gels washed twice with PBS. Finally, 3x 10^4^ cells were seeded on the gels and allowed to attach overnight. Talin deletion was achieved by adding 5mM of 4-OHT (Sigma-Aldrich Cat# H7904) to the culture medium post-spreading. At the endpoint of talin deletion immunostaining with phalloidin was performed to visualize cell shape. The imaging was performed using LEICA Confocal SP8 (WLL) microscope.

### Traction force microscopy

Primary mouse lung endothelial cells (MLECs) were plated on polyacrylamide gels with a stiffness of ∼11.29+-0.49kPa. Briefly, glass-bottom dishes (ibidi) were silanized with 0.5% 3-aminopropyltrimethoxy-silane (ACROS), and activated with 0.5% glutaraldehyde (Sigma). A drop of 10ul of gel solution containing acrylamide (7.49%), bis-acrylamide (0.1%), ammonium persulfate (APS), TEMED and FluoSpheres® carboxylate-modified microspheres (diameter 0.2 μm, Molecular Probes-F8801) was added to the dishes and covered by a pre-coated with 1mg/ml fibronectin 13mm coverslip. After gel polymerization, the top coverslip was removed 10^5^ cells was seeded on the gels. Following cell attachment and spreading for at least overnight, 5mM 4-OHT (Sigma-Aldrich Cat# H7904) were added to the medium and cells were cultured until the endpoint of talin deletion. Images were acquired before and after release of cells with Trypsin-EDTA using a ZEISS AxioObserver Z.1 microscope and proceeded for analysis. Bead images were aligned to correct experimental drift using ImageJ prior to analysis. Traction force was measured as described previously (*67, 68*). Force reconstruction was conducted with the assumption that the substrate is a linear elastic half space, using Fourier Transform Traction Cytometry. The Tikhonov regularization parameter was set to 5 × 10−20 for all experiments. The position of the beads in reference image and deformed one was measured using MTT algorithm (*131*). The problem of calculating the stress field from the displacement is solved in Fourier space then inverted back to real space. The final stress field is obtained on a grid with 0.432 μm spacing (four pixels). All calculations and image processing were performed in Matlab.

### Western blot analysis

Whole endothelial extracts were prepared from 70-80% confluent cells in lysis buffer (3% SDS, 60 mM sucrose, 65 mM Tris (pH 6.8) containing protease inhibitor cocktail (Roche-11836170001). Protein extracts were sonicated briefly and the concentration of the isolated proteins was assessed using BCA Protein Assay Reagent (Thermo Scientific™, PI23227). The desirable amount of the protein lysate (15 – 30 μg) was denatured and separated in a 3-6% or 5-10% SDS–PAGE gel. Following transfer to polyvinylidene difuoride (PVDF, Millipore-IPVH00010) membrane by electrophoresis, PVDF membranes were blocked for 1hour in 3% or 5% milk in Tris-buffered saline with 0.1% Tween-20 (TBS-T). After blocking, the membrane was incubated with primary antibodies diluted in 3-5% BSA in PBS-T overnight at 4°C. The next day, membranes washed three times with TBS-T, before incubation with the appropriate HRP-conjugated secondary antibodies in blocking buffer at room temperature for 1 hour. Signal development was performed using the LuminataTM Crescendo Western HRP Substrate (Millipore) and ChemiDoc^TM^ XRS+ was used to visualize the bands. Quantification of bands from Western blot was performed with Image Lab^TM^ (Bio-Rad) software. The following antibodies were used for immunoblotting: Talin-1 (1:5000 dilution, Mouse monoclonal, Clone 97H6; Bio-Rad, Cat# MCA4770), YAP (1:1000 dilution, Mouse monoclonal; Santa Cruz, Cat# sc-101199), VE-cadherin (1:1000 dilution, BD Biosciences, Cat# 555289), actin (1:2000 dilution, Rabbit polyclonal; Sigma-Aldrich, Cat# A2103), phospho-myosin light chain (1:1000 dilution, Rabbit polyclonal; Cell Signaling, Cat# 3671), b1-integrin (1:1000 dilution, Rat monoclonal, clone MB1.2; Millipore, #MAB1997), HSC70 (1:5000 dilution, Mouse monoclonal; Santa Cruz, #sc7298), GAPDH (1:5000 dilution, Mouse monoclonal, clone 6C5; Millipore, #MAB374). Uncropped versions of the Western blots are provided in supplementary figure 14.

### RNA isolation and Quantitative real-time PCR

Primary endothelial cells were lysed in RNeasy Lysis buffer (Qiagen, 79216). Total mRNA was extracted from primary cells using the NucleoSpin^R^ RNA (Macherey Nagel, 740955.10) and sample RNA concentrations were quantified using a Nanodrop ND-10000 spectrophotometer. 1-3 μg of total RNA was reverse-transcribed to cDNA using RevertAid First Strand cDNA Synthesis Kit (Thermo Fisher Scientific K1621) according to the manufacturer instructions. Quantitative real-time PCR was performed with StepOne Plus Real-Time PCR system (Applied Biosystems) using SYBR Green PCR Master Mix (Thermo Fisher Scientific, 4472920). Relative gene expression of *TLN1* and known YAP target genes (*Ankrd1* & *Cyr61*) was analyzed by the DCt method normalized to *RPLP1* mRNA. The primers used were: mouse *RPLP1*, forward: 5’-GTG GAG GCA AAG AAG GAA GA-3’ and reverse: 5’-TCA GCT CTT TAT TAG CCA ACT TAA C-3’; mouse *TLN-1*, forward: 5’-TCG CTG AAG ATT AGC ATT GG-3’ and reverse: 5’-GGG TCA TCA TCT GAC AGA AAG-3’; mouse *Ankrd1*, forward: 5’-GCT GGT AAC AGG CAA AAA GAA C-3’ and reverse: 5’-CCT CTC GCA GTT TCT CGC T-3’; mouse *Cyr61*, forward: 5’-CTG CGC TAA ACA ACT CAA CGA-3’ and reverse: 5’-GCA GAT CCCTTT CAG AGC GG-3’. Analysis was performed using StepOne Software v2.3.

### Statistical analysis

Experiments were repeated minimum three times or as required for the effect size or sample variation, especially for animal experiments. Graphs were designed and analysed using Prism Software 8.0 (GraphPad, La Jolla, CA, USA). All data from *in vivo*, *ex vivo* and *in vitro* experiments are presented as mean ± standard error of the mean (s.e.m) or violin plots. Statistical analysis was performed using non-parametric Mann-Whitney *U* rank sum test (*P<0.05, **P<0.01, ***P<0.001 and ****P<0.0001, ns, non-significant). More details about the sample size used in each experiment and the p values derived from the statistical analysis are presented in figure legends. The Materials and Methods section should provide sufficient information to allow replication of the results. Begin with a section titled Experimental Design describing the objectives and design of the study as well as prespecified components.

In addition, include a section titled Statistical Analysis at the end that fully describes the statistical methods with enough detail to enable a knowledgeable reader with access to the original data to verify the results. The values for *N*, *P*, and the specific statistical test performed for each experiment should be included in the appropriate figure legend or main text.

All descriptions of materials and methods should be included after the Discussion. This section should be broken up by short subheadings. Under exceptional circumstances, when a particularly lengthy description is required, a portion of the Materials and Methods can be included in the Supplementary Materials.

## Acknowledgments

The authors would like to thank BSRC Fleming animal facility, and BioImaging facility, Prof. David Critchley (University of Leicester) for providing the talin floxed mice and critically reading the manuscript, Prof. Marcus Fruttiger for providing the Pdgfb-iCreER^T2^ mice, Kostourou Lab members, Dr George Panayotou, Dr Christos Zervas and Prof. Kairbaan Hodivala-Dilke for insightful comments and helpful suggestions.

## Funding

World Wide Cancer Research (WWCR), grant number 14-1260 Operational Programme “Competitiveness, Entrepreneurship and Innovation” (NSRF 2014-2020) co-financed by Greece and the European Union, Action: Reinforcement of the Research and Innovation Infrastructure, ‘‘A Greek Research Infrastructure for Visualizing and Monitoring Fundamental Biological Processes (BIO-IMAGING-GR)’’ (MIS 5002755) The Hellenic Foundation for Research and Innovation (H.F.R.I.) under the “First Call for H.F.R.I. Research Projects to support Faculty members and Researchers and the procurement of high-cost research equipment grant” (Project Number: HFRI-FM17-2644)

## Author contributions

NP performed experiments and analysed data, CA performed and analysed all retina experiments, GR performed cell transfections and stainings and analysed data, DM analysed TFM data, AG and CD performed and guided μPET/CT imaging, VK conceived the study, designed experiments, performed initial *in vivo* experiments, analysed and interpreted data, acquired funding and wrote the manuscript.

## Competing interests

Authors declare that they have no competing interests.

## Data and materials availability

All data are available in the main text or the supplementary materials. Mice are subject to material transfer agreements (MTAs).

---

**Fig. S1.**
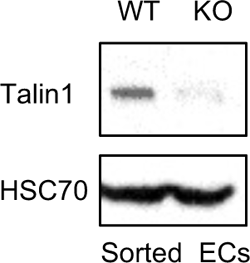
Inducible *in vivo* endothelial talin deletion. Representative western blot analysis of talin protein expression levels in primary endothelial cell extracts isolated from EC-Talin^WT^ and EC-Talin^KO^ mice bearing B16F0 tumours, showing the elimination of talin expression. HSC70 was used as a loading control.

**Fig. S2.**
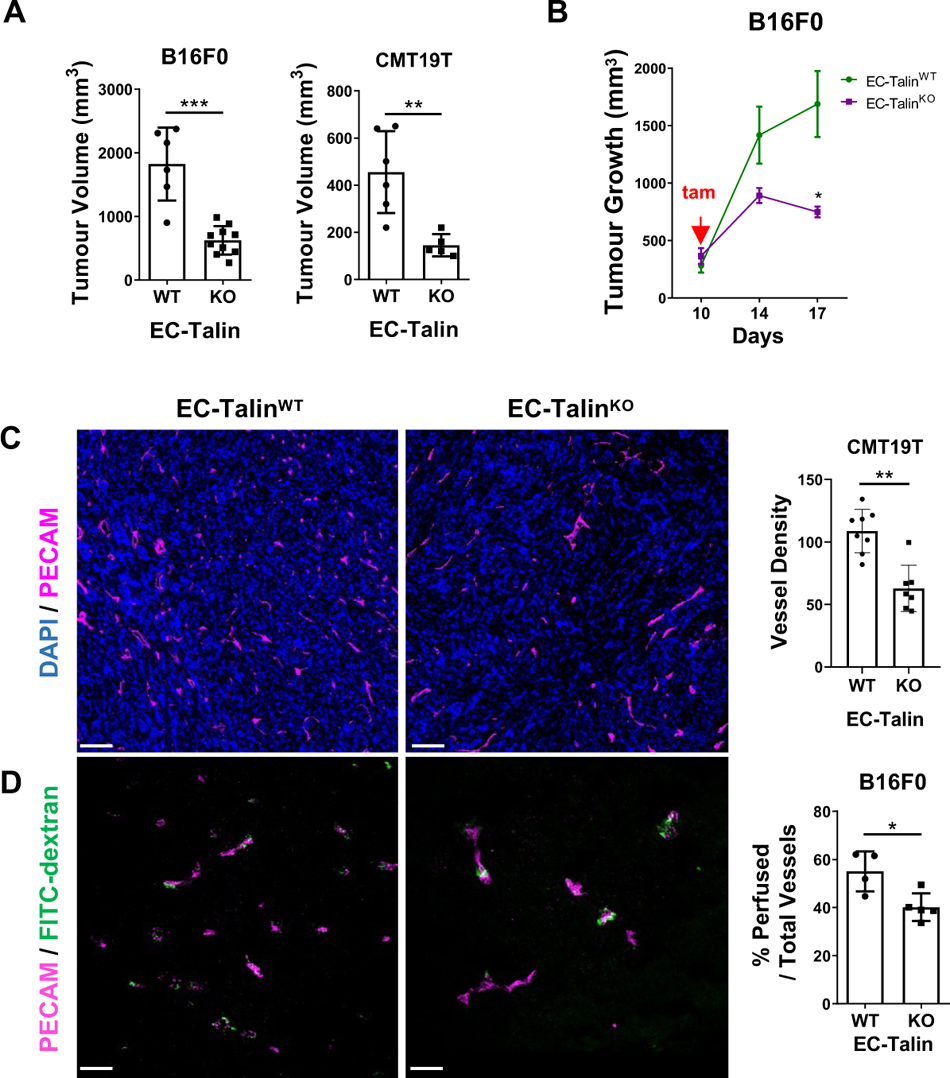
Endothelial loss of talin after tumour initiation limits tumour growth and causes tumour regression. (**A**) Bar charts represent tumour volumes of B16F0 and CMT19T tumours grown in EC-Talin^WT^ and EC-Talin^KO^ mice for 14 days and 16 days respectively. Endothelial deletion of talin was induced by tamoxifen treatment 5 days after subcutaneous tumour cell inoculation. B16F0 studies: n=6 EC-Talin^WT^ and 10 EC-Talin^KO^ mice, ***P=0.0005; CMT19T studies: n=6 EC-Talin^WT^ and 5 EC-Talin^KO^ mice, **P=0.0065. (**B**) Line graph represents the growth curve of tumours after endothelial talin deletion in 10 days-established B16F0s, mean tumour volume ± s.e.m. on day 10, day 14 and day 17; day 10: n=9 EC-Talin^WT^ and 7 EC-Talin^KO^ mice; day 14: n=7 mice per genotype; day 17: n=5 EC-Talin^WT^ and 7 EC-Talin^KO^ mice, day 10 and 14: no statistically significant difference (ns), day 17: *P=0.0101. (**C**) Representative images of 10μm CMT19T tumour sections from EC-Talin^WT^ and EC-Talin^KO^ mice stained with antibodies against PECAM to demarcate blood vessels, displaying reduced tumour angiogenesis upon endothelial loss of talin. DAPI depicts nuclei. Vessel density is given as the number of blood vessels per mm^2^ area of midline tumour section; n=8 EC-Talin^WT^ and 7 EC-Talin^KO^ mice, **P=0.0012. **d** Representative images of 10μm B16F0 tumour sections showing reduced tumour vessel perfusion after intravenous injection of FITC-Dextran in EC-Talin^WT^ and EC-Talin^KO^ mice. Blood vessels were immunodetected with PECAM antibody. Bar chart represents the percentage of the number of perfused vessels over total blood vessels per midline tumour section; n=4 mice per genotype, *P=0.0317. Scale bars, 100 µm (**C**); 50 µm (**D**). All grafs represent the mean value ± s.e.m.; Statistical analysis, Mann–Whitney rank-sum test.

**Fig. S3.**
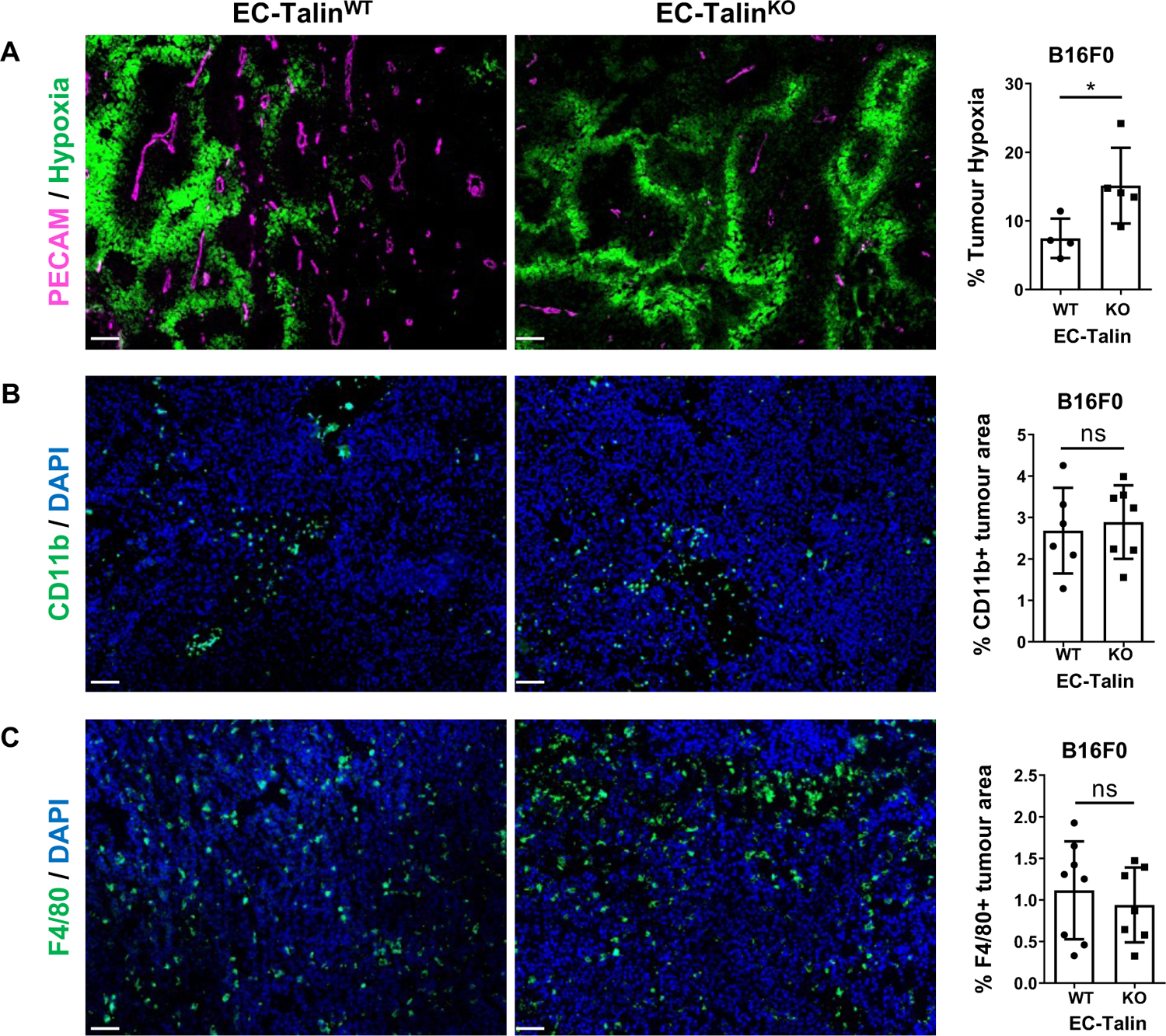
Endothelial loss of talin increases tumour hypoxia without altering tumour immune cell infiltration. (**A**) Endpoint tumour hypoxia was elevated in B16F0 tumours grown in EC-Talin^KO^ mice compared to EC-Talin^WT^ mice. Bar chart represents the percentage of the hypoxic area over the total area of midline tumour sections; n=4 EC-Talin^WT^ and 5 EC-Talin^KO^ mice, *P=0.0317. (**B**) Representative images of 10mm B16F0 tumour sections from EC-Talin^WT^ and EC-Talin^KO^ mice stained with antibodies against CD11b showed no changes in monocyte immune cell infiltration. Bar charts represent the percentage of CD11b positive area over the total tumour area; n=6 EC-Talin^WT^ and 7 EC-Talin^KO^ mice, no statistically significant difference (ns). (**C**) Representative images of 10μm B16F0 tumour sections from EC-Talin^WT^ and EC-Talin^KO^ mice stained with antibodies against F4/80 showed no alteration in macrophage infiltration. Bar charts represent the percentage of F4/80 positive area over the total tumour area; n=8 EC-Talin^WT^ and 7 EC-Talin^KO^ mice, ns. Scale bars, 100 μm. All bar charts represent the mean value ± s.e.m.; Statistical analysis, Mann–Whitney rank-sum test.

**Fig. S4.**
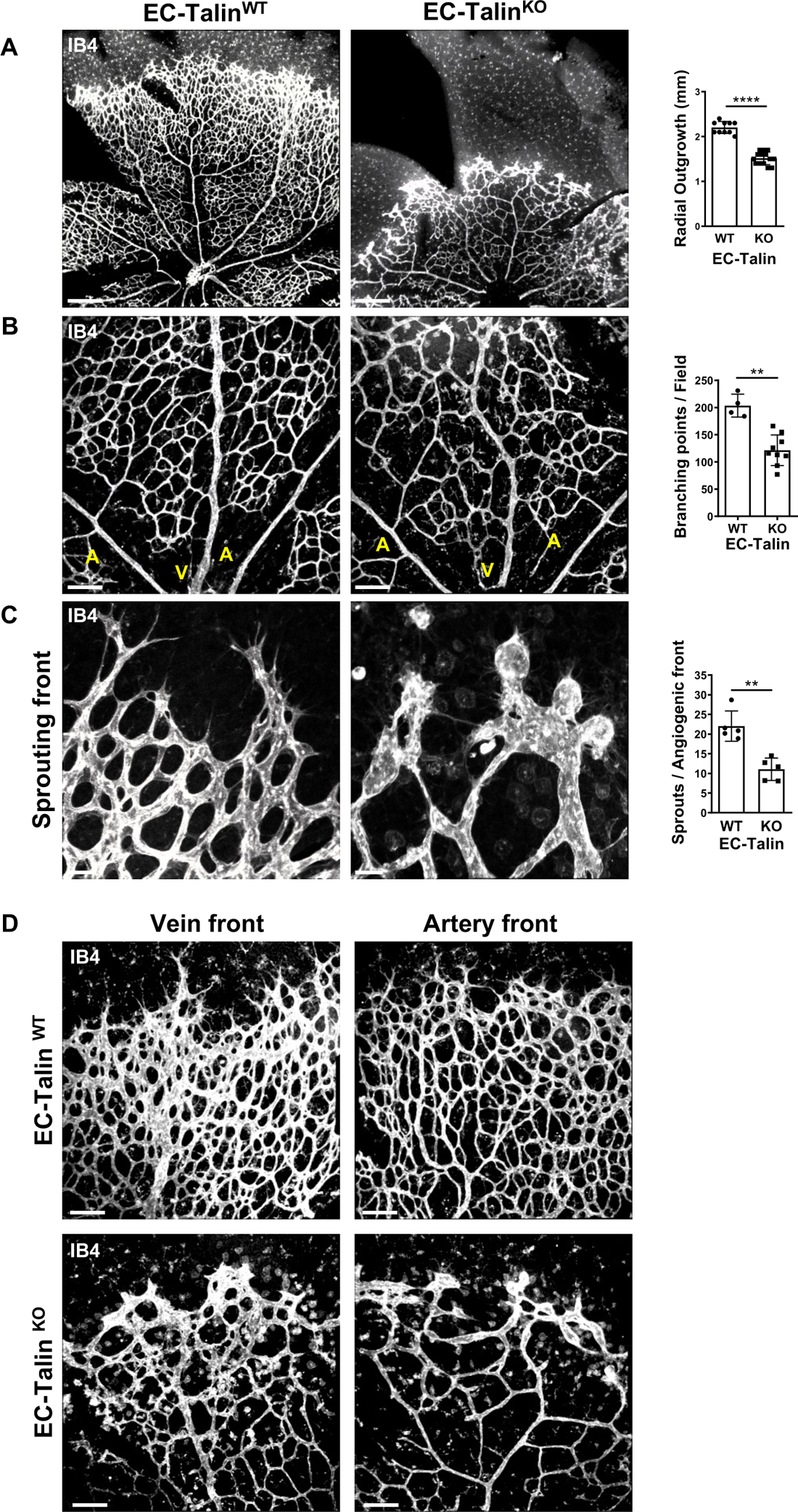
Endothelial loss of talin disrupts retina vascularization at postnatal day 6. (**A**) Low magnification view of the angiogenic front from isolectin B4 (IB4) stained P6 retina showing reduced vessel expansion to the periphery in EC-Talin^KO^ mice compared to EC-Talin^WT^ littermates. Bar chart represents the length of the radial vascular outgrowth from the optic nerve; n=10 EC-Talin^WT^ and 14 EC-Talin^KO^ retinas, ****P<0.0001. (**B**) Confocal images of IB4 stained blood vessels in P6 retina whole mounts, depicting a less dense inner plexus of the developing vasculature. V and A denotes veins and arteries, respectively. Bar chart represents the average number of branching points in randomly selected fields per retina; n=4 EC-Talin^WT^ and 9 EC-Talin^KO^ retinas, **P=0.0028. (**C**) High magnification of the angiogenic front from IB4 stained P6 retinas showing blind ends and EC bulges in EC-Talin^KO^ mice compared to EC-Talin^WT^ littermates. Bar chart represents the number of sprouts per angiogenic front; n=4 EC-Talin^WT^ and 5 EC-Talin^KO^ retinas, **P=0.0079. (**D**) Confocal images of IB4 stained P6 retinas showing reduced vessel density and deformed endothelial sprouts in both arterial and venous angiogenic front in EC-Talin^KO^ mice compared to EC-Talin^WT^ littermates. Experimental observations from at least n=6 retinas per genotype. Scale bars, 300 μm (**A**); 100 μm (**B,D**); 30 μm (**C**)

**Fig. S5.**
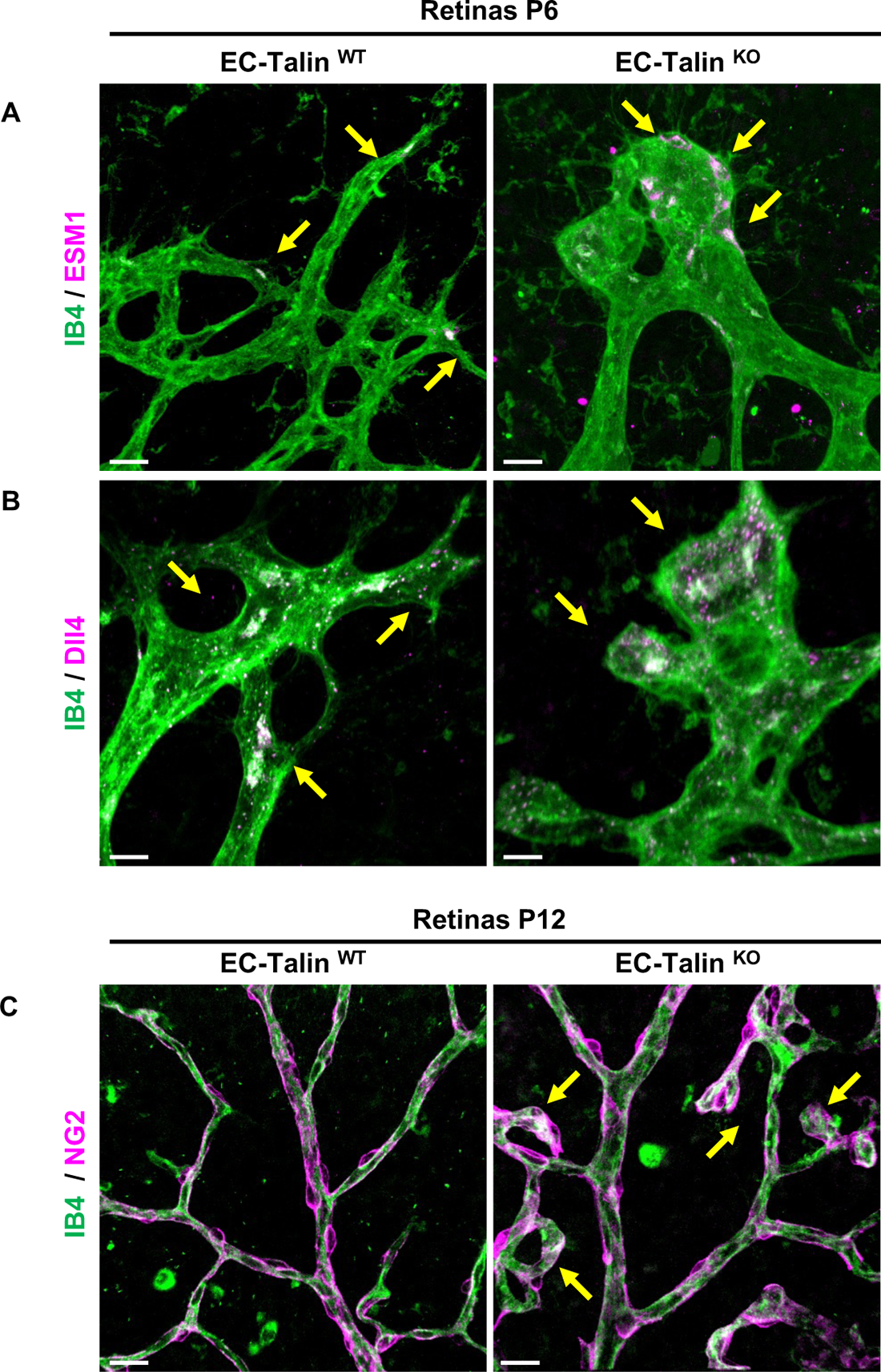
Endothelial deletion of talin does not affect tip-stalk cell selection and pericyte coverage. (**A-B**) High magnification view of angiogenic sprouts in P6 retina whole-mounts stained with isolectin B4 (IB4) and antibodies against (**A**) ESM or (**B**) Dll4 EC tip markers, displayed normal tip-stalk EC differentiation. (**C**) Confocal images of the inner vascular plexus stained with IB4 and pericytes immunodetected by NG2 in P12 superficial layer from EC-Talin^WT^ and EC-Talin^KO^ retinas. Note the NG2 positive pericytes surrounding deformed EC bulges (arrowheads) in EC-Talin^KO^ retinas. Scale bars, 20 μm (**A,C**); 10 μm (**B**). Experimental observations from at least n=3 retinas per genotype.

**Fig. S6.**
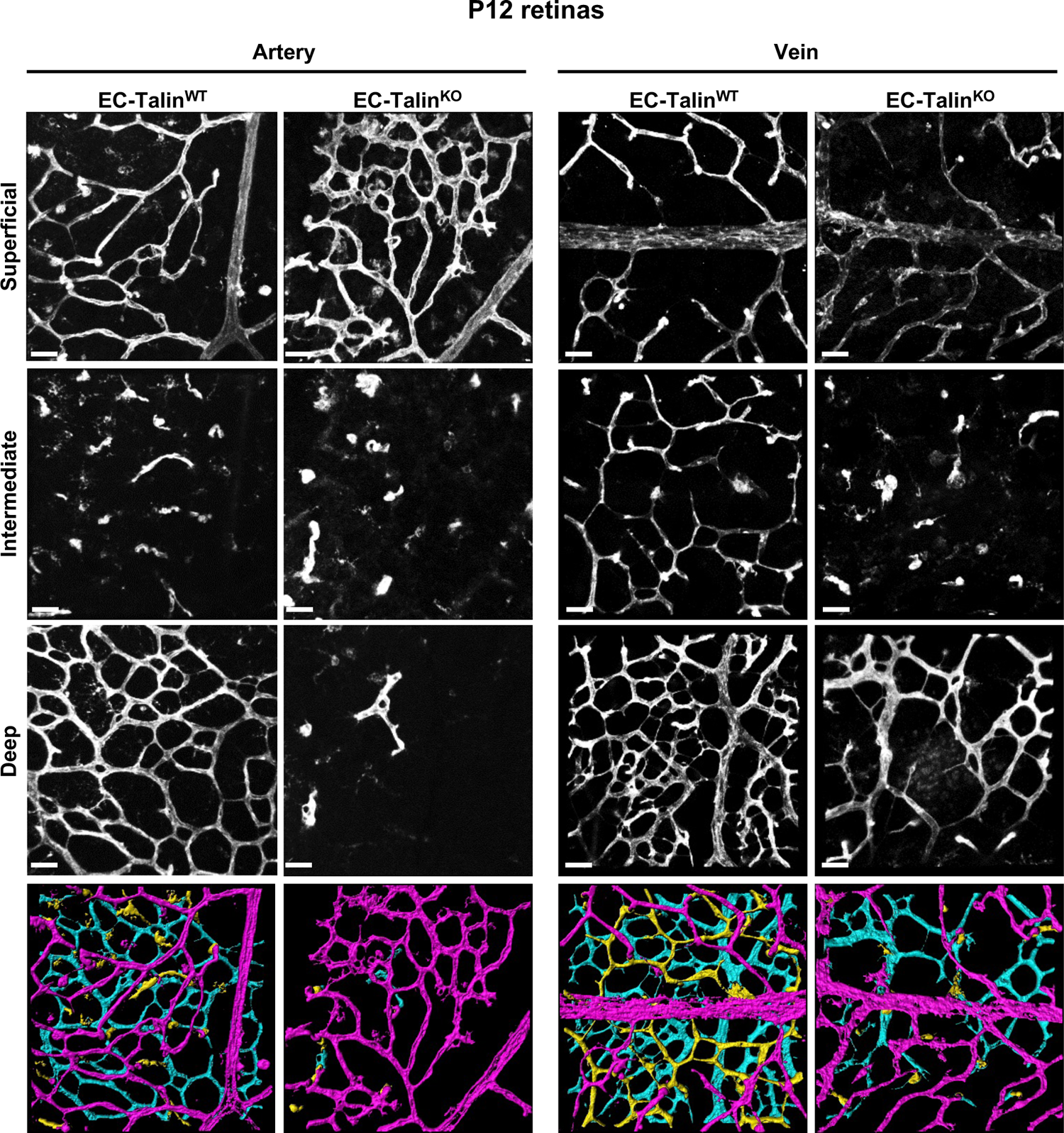
Endothelial talin deletion abolishes tip cell invasion and vascular plexus formation in the deeper retina at postnatal day 12. Confocal images of whole-mount retinas stained for isolectin B4 (IB4) depicting the superficial, intermediate, and deep layers and a pseudo-coloured overlayed image of all layers. Color depth projections show superficial, intermediate, and deep layer as magenta, cyan and yellow, respectively. Talin deletion was induced by 4-OHT administration on postnatal day 6, 7, 8 and retinas were examined on day 12. Both arterial and venous fields displayed defective deep layer formation and lack of intermediate plexus in EC-Talin^KO^ mice compared to EC-Talin^WT^ littermates. Scale bars, 30 μm. Experimental observations from at least n=6 retinas per genotype.

**Fig. S7.**
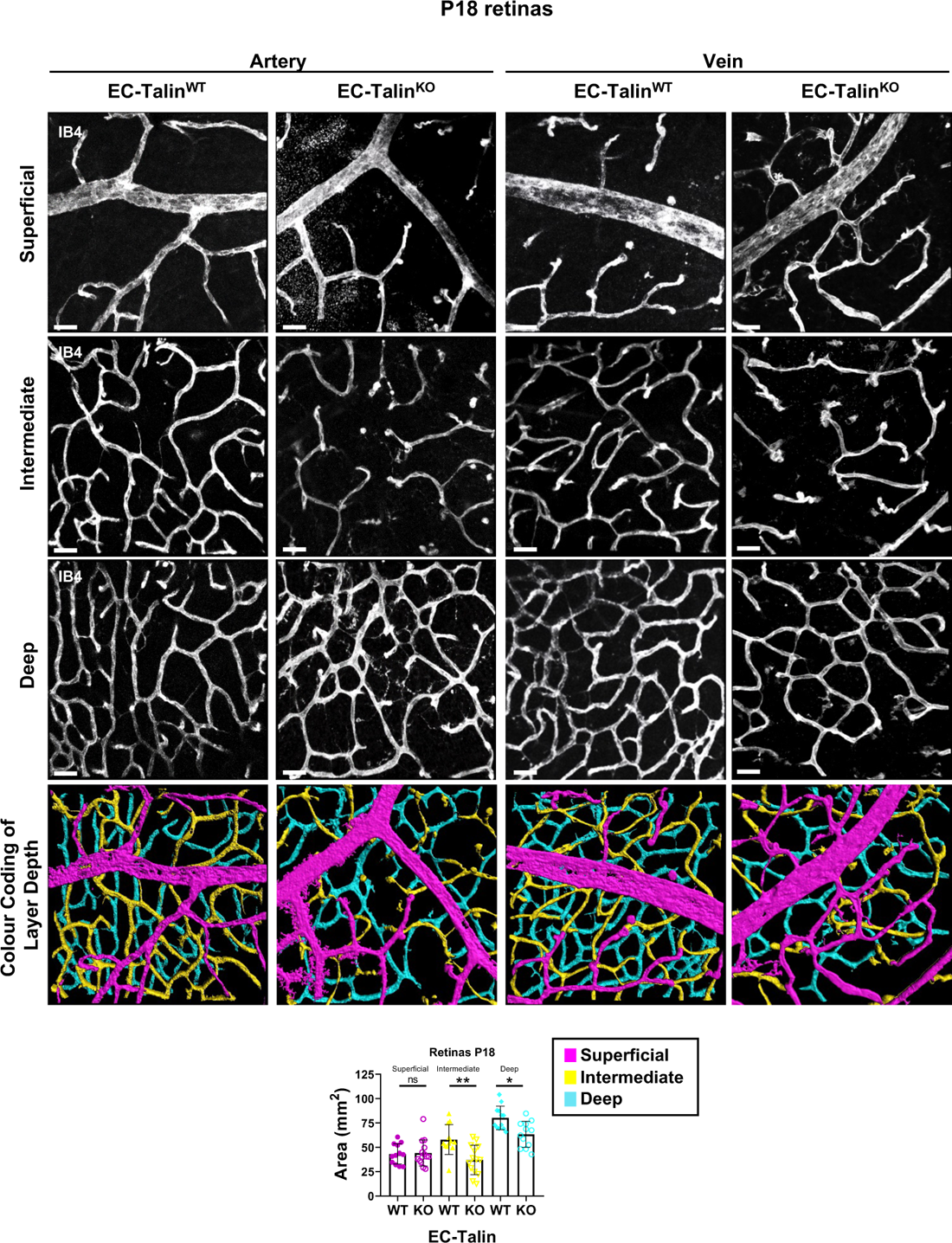
Endothelial loss of talin impairs the growth and compromises the stability of newly formed vascular network at P18 retinas. Confocal images of whole-mount retinas stained for isolectin B4 (IB4) depicting the superficial, intermediate, and deep layers and a pseudo-coloured overlayed image of all layers. Color depth projections show superficial, intermediate, and deep layer as magenta, cyan and yellow, respectively. Talin deletion was induced by 4-OHT administration on postnatal day 14, 15, 16 and retinas were examined on day 18. Reduced vascularization of the intermediate layer and disruption of the newly formed deep layer was observed in both arterial and venous fields of EC-Talin^KO^ compared to EC-Talin^WT^ retinas. Bar charts represents the area of IB4-stained vascular network in each layer from random selected fields, mean ± s.e.m.; n=4 EC-Talin^WT^ and n=5 EC-Talin^KO^ retinas, Statistics: Superficial, ns; Intermediate, **P=0.0037; deep, *P=0.01. Mann–Whitney rank-sum test. Scale bars, 30 μm.

**Fig. S8.**
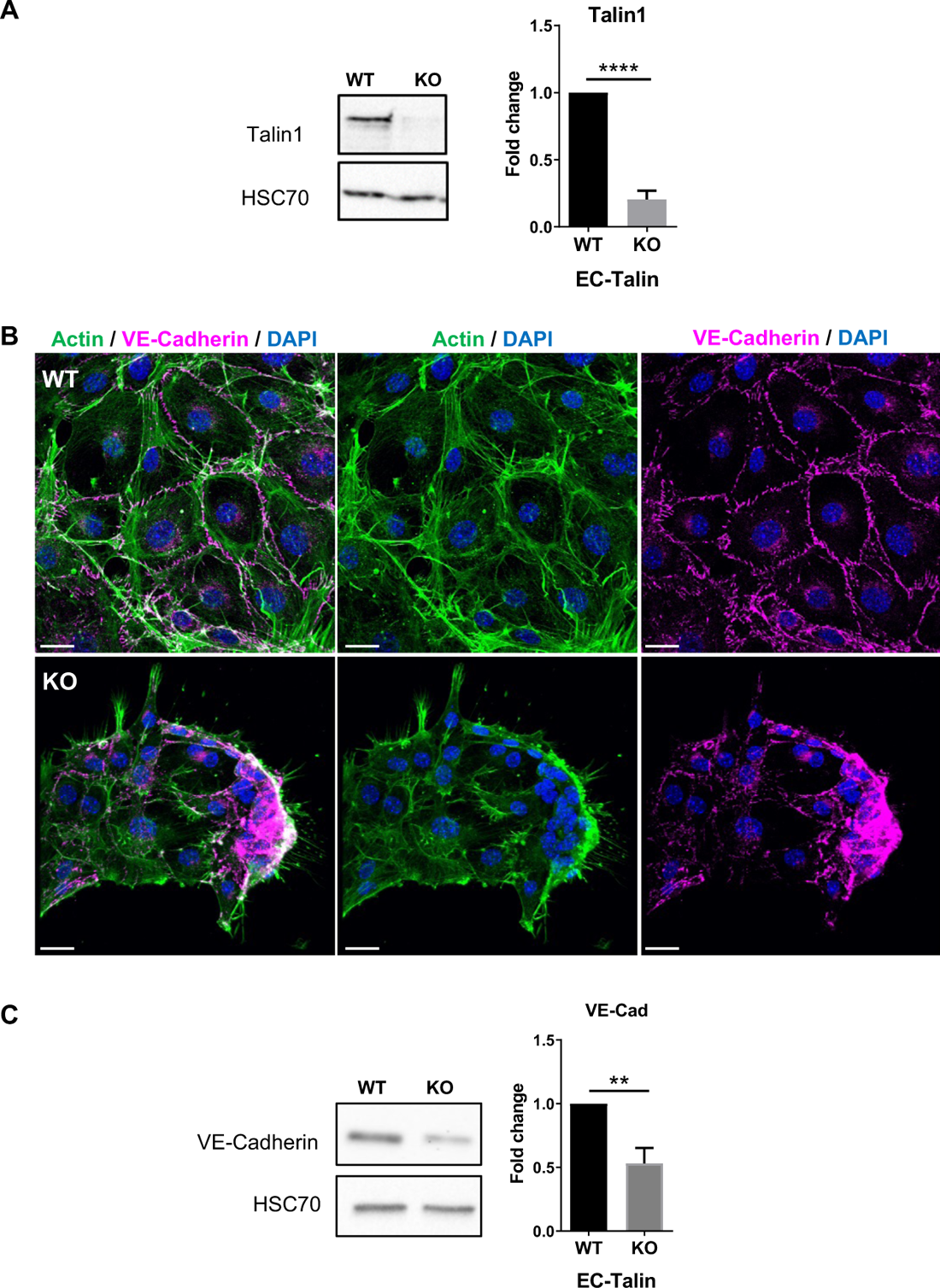
Endothelial loss of talin from confluent monolayers decreases VE-cadherin expression and causes cell aggregation and shrinkage. (**A**) Western blot analysis confirmed significant elimination of talin expression following 4-OHT administration in cell culture of well-adherent primary mouse endothelial cells (KO) compared to control vehicle treated ECs (wt). HSC70 acted as a loading control. Bar chart represents the fold reduction in talin expression, mean ± s.e.m., n=3 independent experiments (wt values set to 1), ****P<0.0001, two-sided Student’s *t*-test. (**B**) Representative confocal images of wt and talin KO endothelial monolayers stained with phalloidin and VE-cadherin, displaying disruption of monolayer organization and cell retraction. DAPI counterstained nuclei. Scale bars, 20 μm. (**C**) Western blot analysis of wt and talin KO cell extracts showed significant reduction of VE-cadherin expression upon talin deletion. HSC70 acted as a loading control. Bar chart represents the fold reduction in VE-cadherin expression, mean ± s.e.m., n=3 independent experiments (WT values set to 1), **P<0.01, two-sided Student’s *t*-test.

**Fig. S9.**
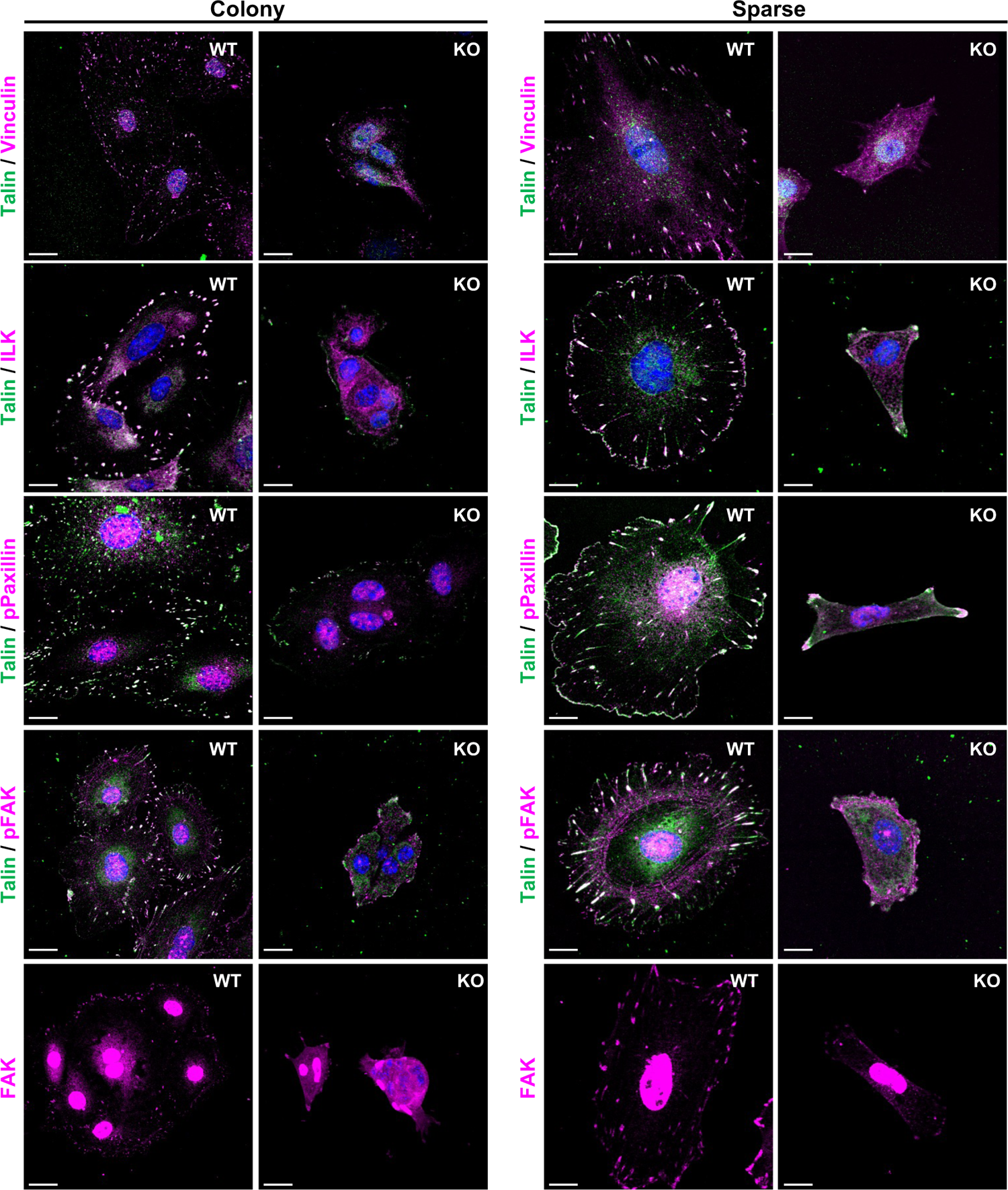
Endothelial talin deletion deconstructs established cell-matrix adhesions in well-adherent cells. Representative confocal images of sparse primary ECs and colonies immunostained with antibodies for talin and key adhesome members. (**A**) talin/vinculin, (**B**) talin/ILK, (**C**) talin/phospho-Y118 paxillin, (**D**) talin/phospho-Y397 FAK and (**E**) total FAK. Nuclei were stained with DAPI. Talin deletion was induced by 4-OHT in well-spread ECs with established cell-matrix adhesions. Note the talin-dependent localization of vinculin, ILK, phospho-paxillin, phospho-FAK and FAK at cell-matrix adhesions. Scale bars, 10 μm (sparse cells); 15 μm (cell colony). Experimental observations from at least n=3 independent experiments with different biological mouse lung EC isolations.

**Fig. S10.**
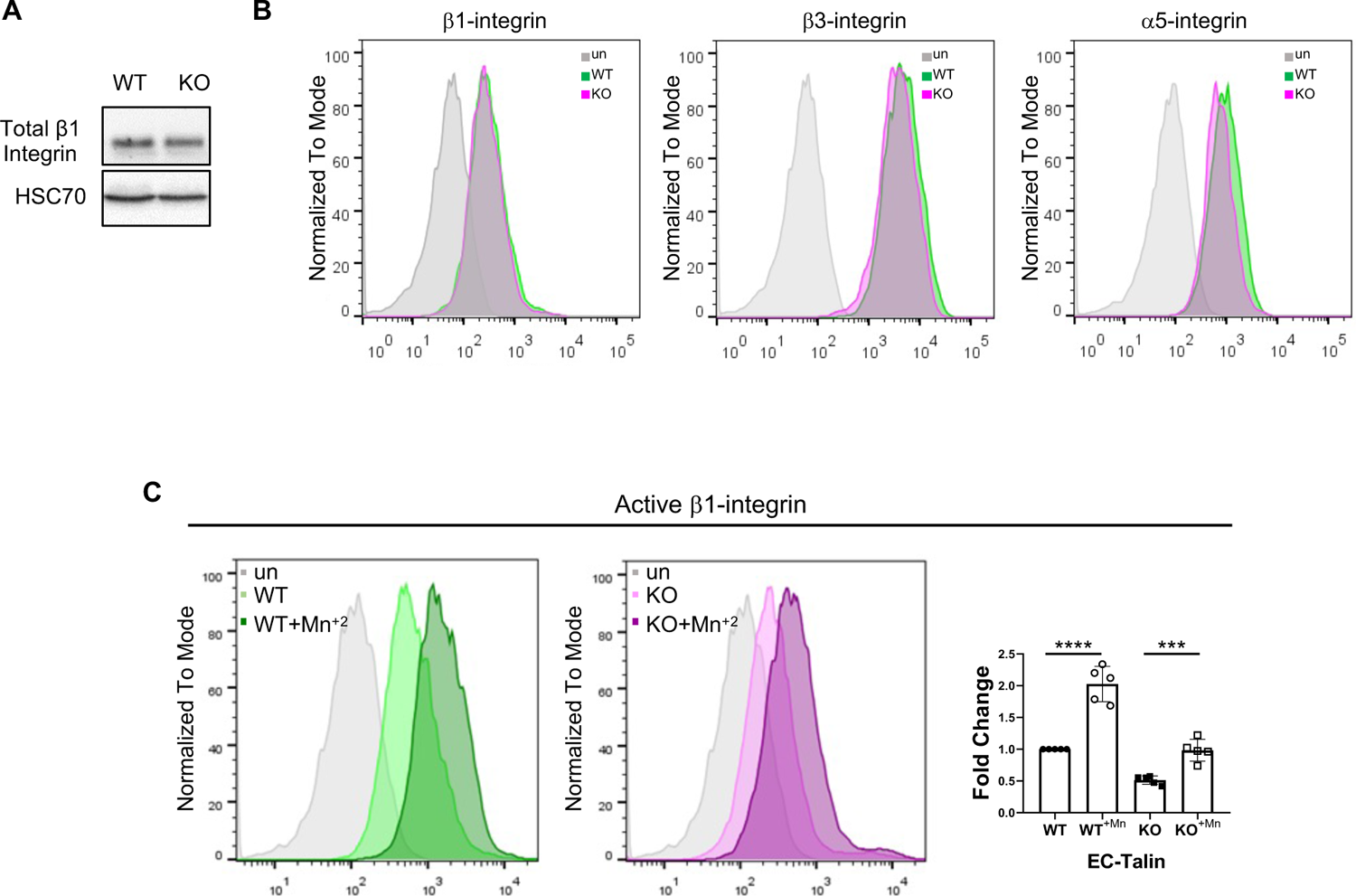
Endothelial loss of talin does not affect surface levels or outside-in activation of blood vessel expressing integrins. (**A**) Representative western blot of b1-integrin expression showed no difference between wt and talin KO EC extracts. HSC70 acted as a loading control. Experimental observation of 3 independent experiments with different biological mouse lung EC isolations. (**B**) FACs histograms of wt (green) and talin KO (purple) ECs immunostained for total surface levels of b1-integrin, b3-integrin and a5-integrin showed no difference between wt and talin KO ECs. (**C**) FACs histograms of wt (green) and talin KO (purple) ECs incubated with or without Mn^2+^ and immunostained with the activation epitope-reporting 9EG7 antibody showed normal Mn^2+^-induced activation of b1-integrin upon loss of talin. Bar chart represents fold changes of active of b1-integrin levels in Mn^2+^ treated and untreated wt and talin KO ECs. Note the 2-fold increase in active b1-integrin levels in both wt and talin KO Mn^2+^ treated ECs. Data are expressed as fold change relative to the wt cells, mean ± s.e.m. from 5 independent experiments. ****P<0.0001 (wt + Mn^+2^ /wt); ***P=0.0004 (KO + Mn^+2^ /KO), two-sided Student’s *t*-test.

**Fig. S11.**
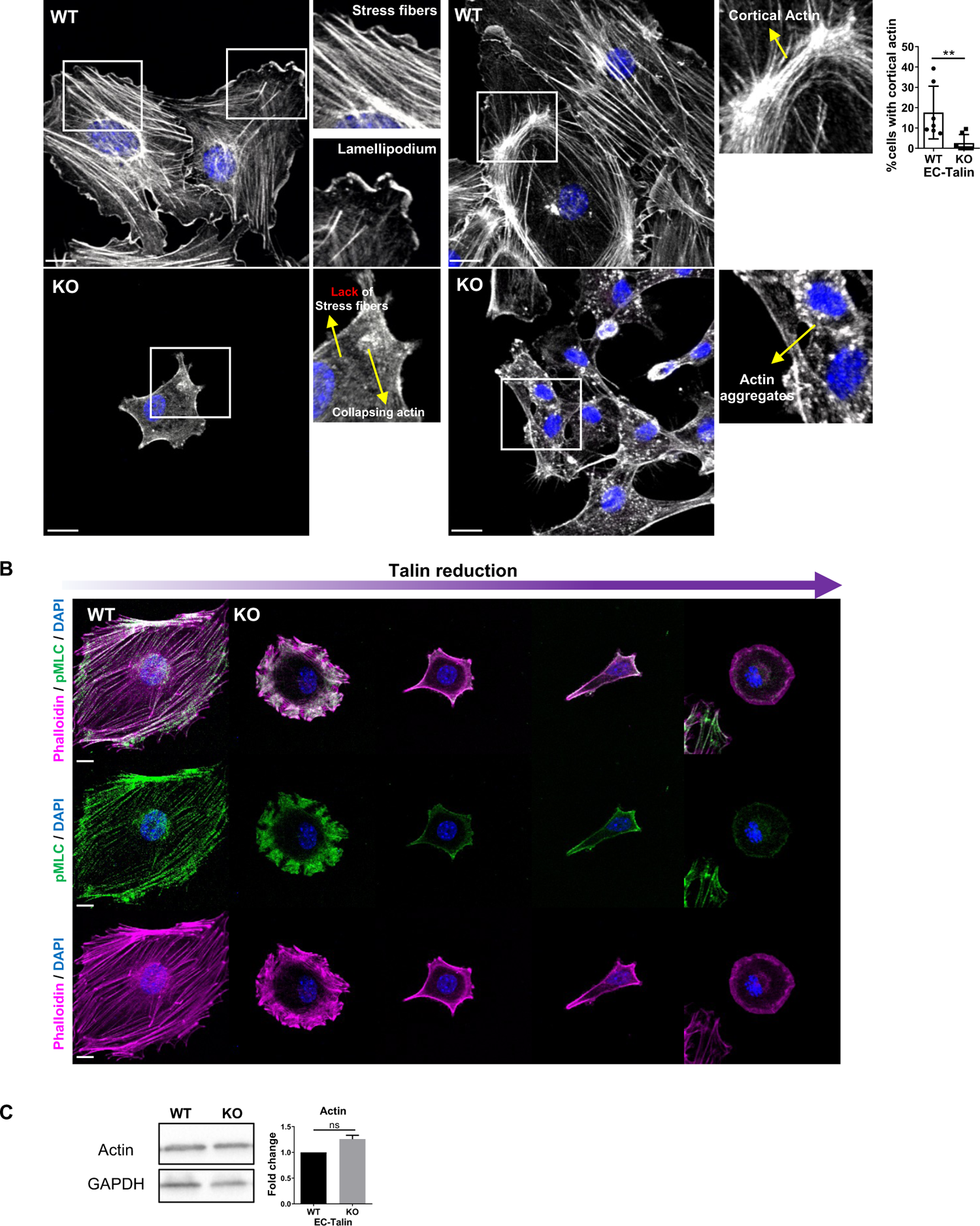
Endothelial loss of talin destroys actin cytoskeletal dynamics in well-adherent cells. (**A**) Confocal images of actin immunodetection in control (wt) and talin KO primary mouse ECs. DAPI was used for nuclei visualization. Insets illustrate lack of stress fibers, lamellipodium and cortical actin and actin aggregates in talin KO ECs. Bar chart represent the percentage of cells containing cortical actin, mean ± s.e.m. n= 7 independent experiments (total cell numbers 380 wt and 183 KO), **P=0.0064 Mann–Whitney rank-sum test. Scale bars, 15 μm. (**B**) Representative confocal images of phalloidin staining to demarcate F-actin and immunodetection of phospho-myosin light chain (pMLC), showed disruption of cytoskeletal organization and contractility upon talin reduction in well-adherent primary mouse ECs. Scale bar, 10 μm. Experimental observations from at least n=3 independent experiments with different biological mouse lung EC isolations. (**C**) Western blot analysis of actin expression in wt and talin KO EC extracts showed no difference. GAPDH acted as a loading control. Bar chart represents the levels of actin expression in wt and talin KO ECs, mean ± s.e.m., n=3 independent experiments (wt values set to 1), Two-sided Student’s *t*-test, ns, no statistically significant difference.

**Fig. S12.**
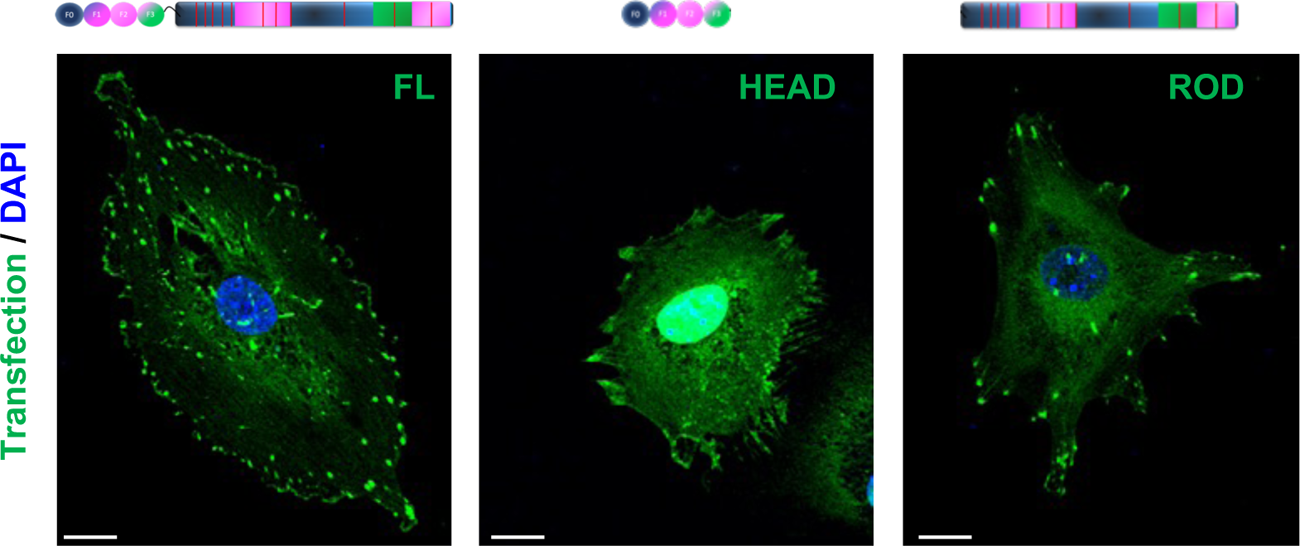
Expression of talin constructs in primary mouse ECs. Representative fluorescent images of primary mouse lung ECs expressing GFP-fused talin constructs, as indicated in schemes, full-length talin (FL, 1-2451aa), talin head domain (HEAD, 1-433aa), talin-rod domain (ROD, 344-2541aa). Nuclei were stained with DAPI. Scale bars, 15 µm.

**Fig. S13.**
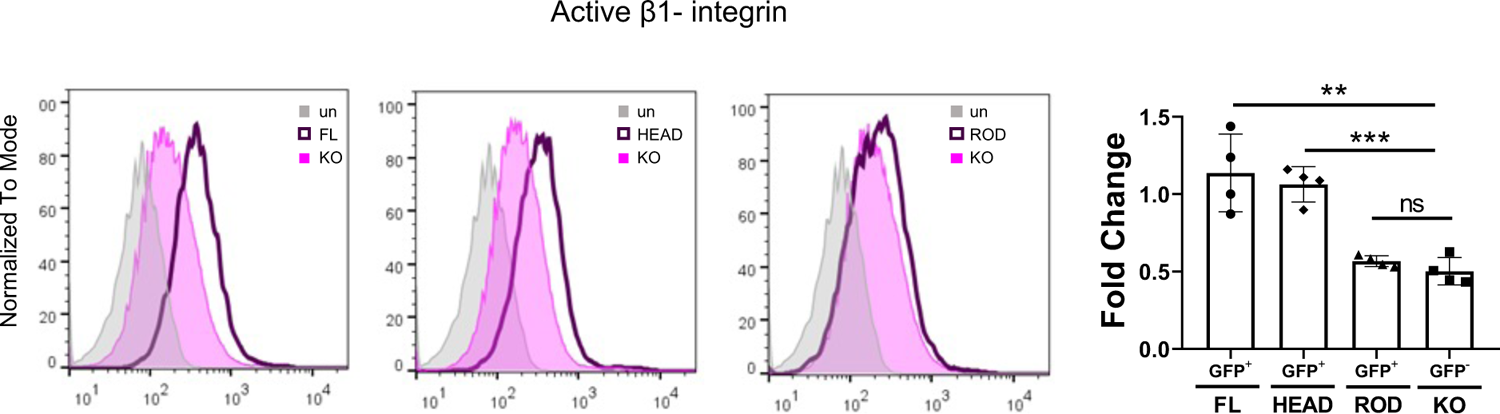
Talin head but not talin rod rescued integrin activation. FACs histograms of talin KO transfected ECs with full-length talin-GFP (FL), talin head-GFP (HEAD) and talin rod-GFP (ROD) stained with the activation epitope-reporting 9EG7 antibody. Primary mouse lung ECs were transiently transfected with the different talin-GFP constructs and treated with 4-OHT to induce talin deletion, generating a mix cell population of transfected GFP positive and untransfected GFP negative talin KO cells. Note that talin head rescued activation of b1-integrin similar to full-length talin, whereas talin rod had no effect. Bar charts represents the fold change in b1-integrin activation in transfected and untransfected talin KO ECs, relative to control cells, mean ± s.e.m, n= at least 4 independent experiments. **P=0.0031 (FL/KO), ***P=0.0002 (HEAD/KO), no significance (ROD/KO), two-sided Student’s *t*-test.

## Notes

### Competing Interest Statement

The authors have declared no competing interest.

